# Genomic signature of kin selection in an ant with obligately sterile workers

**DOI:** 10.1101/072496

**Authors:** Michael R. Warner, Alexander S. Mikheyev, Timothy A. Linksvayer

## Abstract

Kin selection is thought to drive the evolution of cooperation and conflict, but the specific genes and genome-wide patterns shaped by kin selection are unknown. We identified thousands of genes associated with the sterile ant worker caste, the archetype of an altruistic phenotype shaped by kin selection, and then used population and comparative genomic approaches to study patterns of molecular evolution at these genes. Consistent with population genetic theoretical predictions, worker-upregulated genes showed relaxed adaptive evolution compared to genes upregulated in reproductive castes. Worker-upregulated genes included more taxonomically-restricted genes, indicating that the worker caste has recruited more novel genes, yet these genes also showed relaxed selection. Our study identifies a putative genomic signature of kin selection and helps to integrate emerging sociogenomic data with longstanding social evolution theory.

---

Kin selection theory provides the dominant framework for understanding the evolution of diverse types of social behavior, from cooperation to conflict, across the tree of life (Hamilton 1964; Bourke 2011). While kin selection theory has always had an explicit genetic focus (Hamilton 1964), researchers have made little progress in identifying specific genes that have been shaped by kin selection (Thompson et al. 2013; Ronai et al. 2016), or in identifying genome-wide evolutionary signatures of kin selection (Van Dyken and Wade 2012; Ostrowski et al. 2015). This shortfall is particularly notable in the social insects because the sterile worker caste is the archetypical example of an altruistic phenotype that evolved through kin selection (Hamilton 1964; Queller and Strassmann 1998; Bourke 2011).

The caste system of division of labor between reproductive queens and sterile workers, which first evolved in ants over 100 mya (Ward 2014), is a striking evolutionary innovation that enabled the radiation and ecological dominance of insect societies (Hölldobler and Wilson 1990). While queen and worker castes share the same genome, they express alternate suites of derived traits associated with specialization on either reproduction, or on foraging, nest defense, and brood care (Hölldobler and Wilson 1990). Because queens (and their short-lived male mates) reproduce and hence can directly pass their genes to the next generation, their traits are shaped directly by natural selection. In contrast, obligately sterile workers can only pass on their genes indirectly, by helping their fully-fertile relatives to reproduce, so that worker traits are shaped indirectly, by kin selection (Hamilton 1964; Bourke 2011). Population genetic models show that in theory, all-else-equal, genes associated with the expression of worker traits should experience relaxed rates of adaptive molecular evolution compared to genes associated with the expression of reproductive traits, with the degree of relaxation proportional to the relatedness between workers and their fully-fertile relatives (Linksvayer and Wade 2009; Hall and Goodisman 2012; Linksvayer and Wade 2016).

Using the pharaoh ant, *Monomorium pharaonis*, a derived ant with obligately sterile workers and many queens per colony (i.e. low relatedness) (Hölldobler and Wilson 1990), in which signatures of kin selection are expected to be pronounced, we identified caste-associated genes and studied genomic signatures of short- and long-term molecular evolution of these genes. We used a large set of *M. pharaonis* samples (159 total RNA sequencing libraries; Table S1) that included a time series of developing worker and reproductive (i.e. queen and male) larvae as well as adult worker and queen head and abdominal tissue (Fig. 1A) to identify genes that were upregulated in reproductive versus worker castes. The number of differentially-expressed genes between worker and reproductive larvae at each stage increased across larval development, corresponding to divergence for overall body size (Fig. 1B; Fig. S1). Most differentially expressed genes were detected between adult queen and worker abdominal tissue (Fig. 1B), which is expected given that queens have well-developed ovaries in their abdomens while workers lack reproductive organs.

**Fig. 1.**
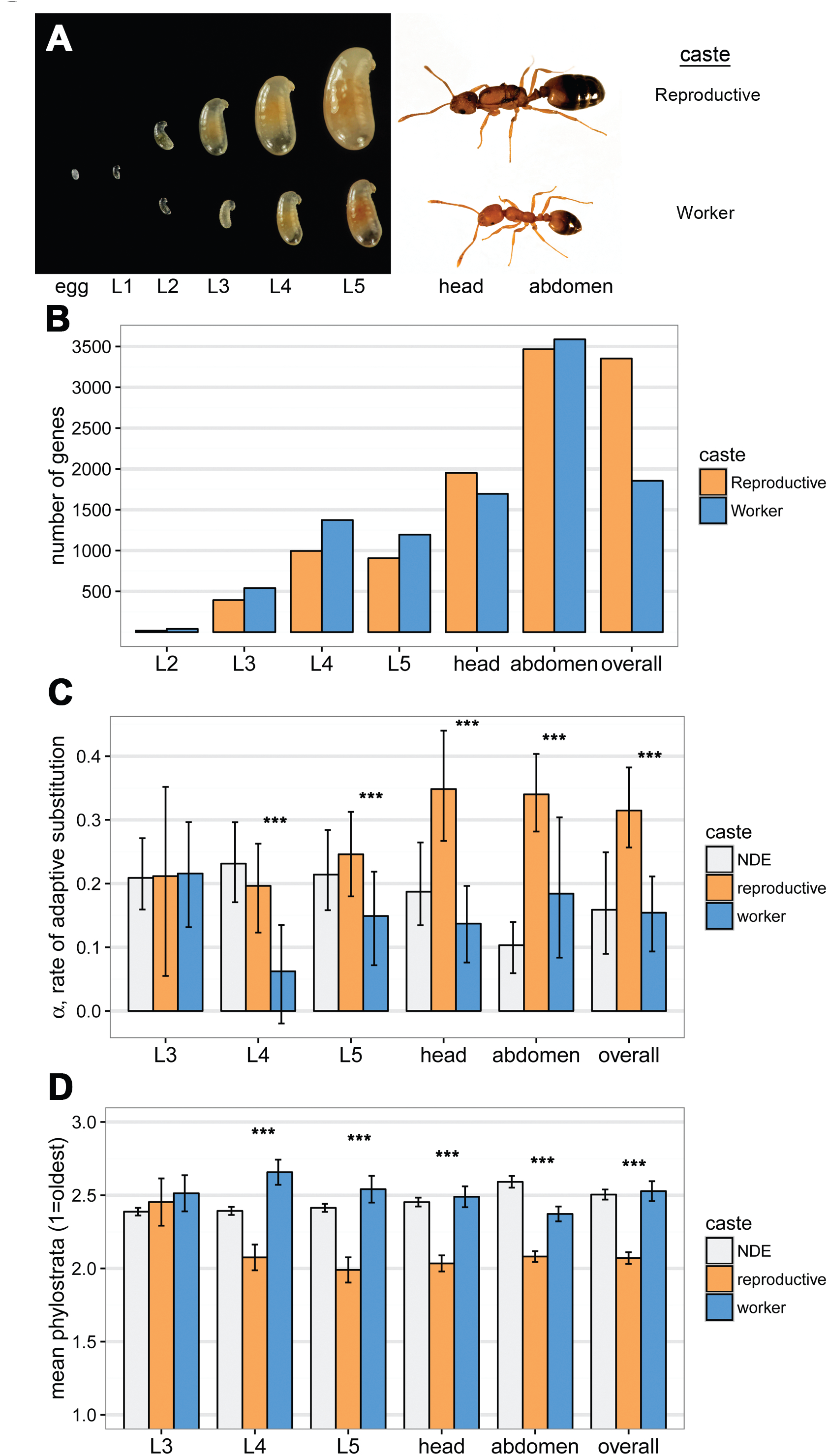
Genomic signature of kin selection. **(A)** In order to identify genes upregulated in reproductives versus worker castes, which should be shaped mainly by direct versus indirect (i.e. kin) selection, respectively, we collected a time series of worker- and reproductive (i.e. queen and male) larvae (L2-L5), as well as adult worker and queen head and abdomen tissue samples. **(B)** Dozens to thousands of genes were differentially expressed and upregulated in either reproductive (orange) or worker (blue) castes for each larval stage and adult tissue sample. “Overall” shows genes that were differentially expressed across all samples (i.e. genes with a main effect of caste on expression). The L2 comparison is excluded from subsequent analyses because only 59 total genes were differentially expressed at this early stage. **(C)** Reproductive-upregulated genes had higher α, the proportion of amino acid substitutions fixed by positive selection, for all comparisons except for L3. NDE, non-differentially expressed genes. **(D)** Reproductiveupregulated genes were also older on average (i.e. lower mean phylostrata) for all comparisons except L3. The phylostrata were grouped into the six categories as shown in Fig. 2A, but using all original 19 categories produced the same result (Fig. S7). *** p<0.001.

Next, to compare rates of adaptive molecular evolution at the identified worker- and reproductive-associated genes, we used a population genomic dataset based on 22 resequenced *M. pharaonis* worker genomes together with a single *M. chinense* worker genome as an outgroup. We estimated α, the proportion of amino acid substitutions fixed by positive selection (Bierne and Eyre-Walker 2004; Welch 2006; Obbard et al. 2009). This proportion for worker-associated genes (0.15, 95% CI 0.09-0.21) is approximately half that of reproductive-associated genes (0.31, 95% CI 0.26-0.38; bootstrap p < 0.001; Fig. 1C), indicating a relaxed rate of adaptive evolution for worker-associated genes. Estimates of mean selection coefficients and selective constraint (Fig. S3) also supported the conclusion that worker-associated genes have experienced relaxed selection compared to reproductive-associated genes. These results are consistent with theoretical expectations (Linksvayer and Wade 2009; Linksvayer and Wade 2016), providing a putative genomic signature of kin selection.

To further elucidate the evolution and genomic basis of ant caste, we used a comparative genomic approach, phylostratigraphy (Domazet-Loso et al. 2007), which estimates the evolutionary age of genes based on whether orthologs can be identified across different strata of the tree of life (i.e. phylostrata). Most of the identified worker- and reproductive-associated genes were ancient (i.e., shared across all cellular organisms, eukaryotes, or bilaterian animals; Fig. 2A), arising long before the evolution of eusociality, consistent with most non-differentially expressed genes in the *M. pharaonis* genome (Fig. 2A) as well as most genes in better annotated insect genomes (Fig. S6). Thus, the evolutionary origin and elaboration of ant caste seems to largely involve the recruitment of ancient genes, as proposed in a series of hypotheses (West-Eberhard 1996; Amdam et al. 2004; Linksvayer and Wade 2005; Amdam et al. 2006; Toth and Robinson 2007). Reproductive-associated genes, which are mainly composed of genes upregulated in adult queen tissues (Fig. S3), were especially enriched for ancient phylostrata, indicating that the evolution of the queen caste mainly involved the recruitment and long-term conservation of ancient genes involved in cellular functions (Tables S6, S7). In contrast, worker-associated genes were younger on average than reproductive-associated genes (Figs. 1D, S7)(glm, z=10.3, df = 12622, p < 0.001), with a relatively larger proportion of genes in younger phylostrata (Figs. 2A, S6; see also (Johnson and Tsutsui 2011; Feldmeyer et al. 2014; Harpur et al. 2014)). Interestingly, worker-associated genes in the youngest phylostrata (hymenopteran- and ant-specific genes) were enriched for chemosensory Gene Ontology categories (Table S7). These genes could putatively underlie ant-specific chemosensory adaptations, however this youngest category of worker-associated genes had α estimates that were not greater than zero (bootstrap p = 0.86; Fig. 2B), indicating that positive selection is not driving molecular evolution at these genes. Thus, the phylostratigraphy results provide further evidence that worker-associated genes experience relaxed selection relative to reproductive-associated genes.

**Fig. 2.**
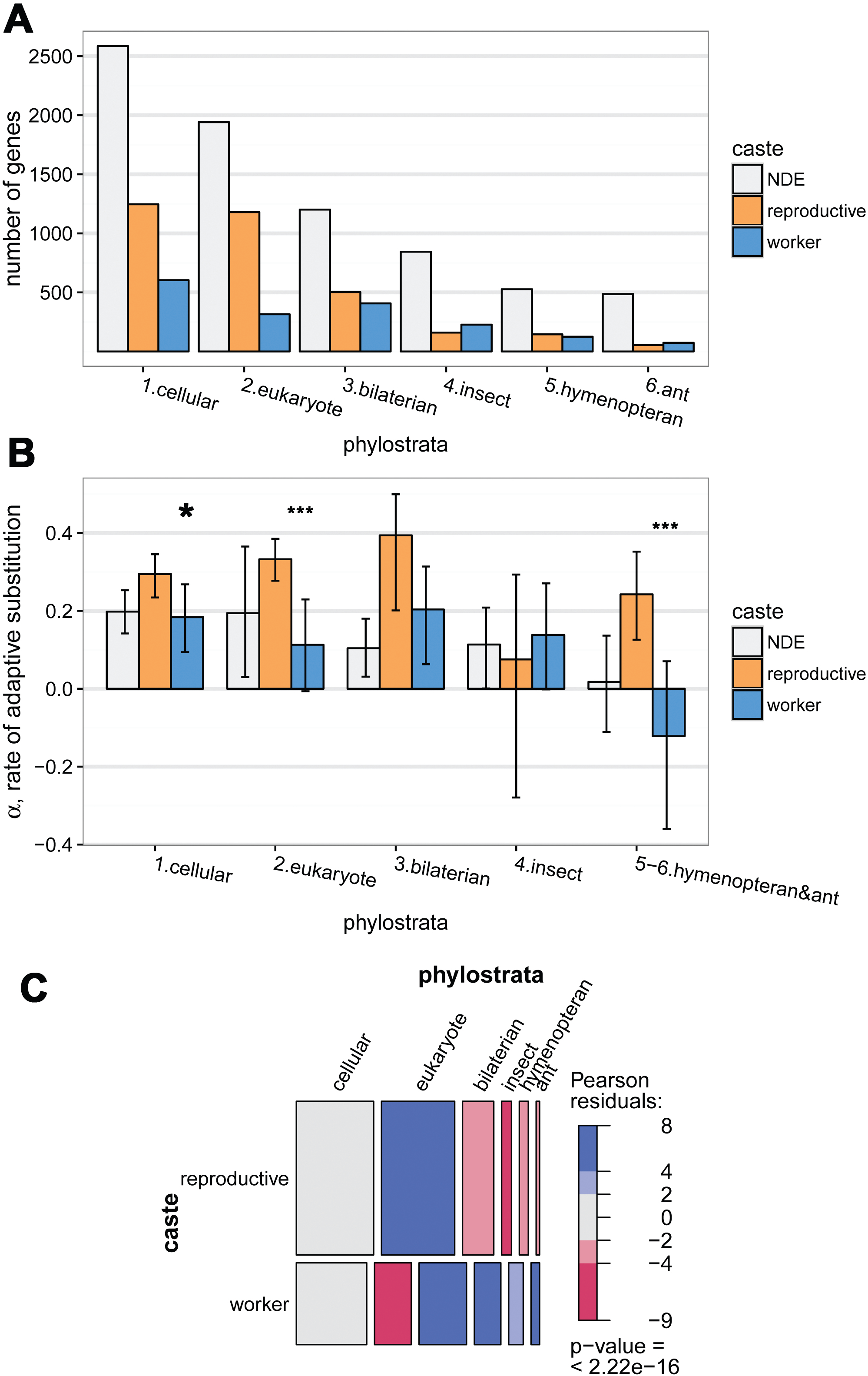
The contribution of ancient and young genes to caste evolution. **(A)** Most caste-associated genes, as well as NDE genes in the *M. pharaonis genome*, are from ancient phylostrata (Fig. S6). **(B)** Genes in the youngest phylostrata tend to show relaxed adaptive evolution (i.e. 95% CI of α overlapping zero), except for reproductive-associated genes in the hymenopteran & ant phylostratum. Genes in the two oldest phylostrata mainly drive the pattern of higher rates of adaptive evolution for reproductive-associated genes relative to worker-associated genes (Fig. 1C); * p<0.05, *** p<0.001. Note that negative α values are caused by sampling error or the presence of mildly deleterious mutations that segregate but do not fix (Obbard et al. 2009). The last two phylostrata (hymenopteran and ant) were combined because there are not enough ant-specific genes for accurate α estimates. **(C)** Mosaic plot showing that relative to reproductive-associated genes, worker-associated genes are enriched for the four youngest phylostrata, while reproductive-upregulated genes are enriched for the eukaryote phylostratum. The area of each cell is proportional to the number of genes in each caste and phylostrata category. Blue shading indicates overrepresentation (light blue p < 0.05, dark blue p < 0.001), and red-shading indicates underrepresentation (light red p < 0.05, dark red, p < 0.001), based on cell standardized pearson residuals.

Recent comparative genomic studies in bees and ants have found signatures of neutral evolution in social insect genomes, thought to be associated with reduced effective population size in species with large societies compared to solitary species (Romiguier et al. 2014; Kapheim et al. 2015). Consistent with these previous findings, our genome-wide estimate of α (0.21, 95% CI 0.17-0.27; Table S3) is lower than most previous estimates from solitary insects such as *Drosophila* (∼0.5) (Bierne and Eyre-Walker 2004; Welch 2006; Obbard et al. 2009; Keightley et al. 2016). Our results indicate that the relatively low genome-wide adaptive substitution rates are at least in part a result of relaxed selection on worker-associated genes, so that neutral evolutionary processes are likely to be especially important for worker-associated genes. It is commonly assumed that intraspecific and interspecific variation for worker morphology and behavior is adaptive (Hölldobler and Wilson 1990; Ferster et al. 2006; Pie and Traniello 2007), but our results suggest that relaxed selection and nonadaptive evolutionary forces also play important roles in the evolution of worker traits, in particular for species with low nestmate relatedness (e.g., due to multiple queens or multiple mating) (Helanterä et al. 2009).

The relaxed selection we observed at worker-associated genes relative to reproductive-associated genes may also result from actual relaxed phenotypic selection on worker traits, in addition to being caused by the fact that worker-associated genes experience mainly indirect selection (Linksvayer and Wade 2009; Linksvayer and Wade 2016). For example, strong phenotypic selection on reproductive traits, caused by intersesexual conflict, is commonly thought to drive elevated rates of adaptive molecular evolution at genes with reproductive function (e.g., seminal proteins) in solitary organisms (Pröschel et al. 2006; Ellegren and Parsch 2007). Whether this pattern could also be true for social insects, where both reproductive and non-reproductive castes are essential for colony survival and reproduction, is not clear, and more research into the relative magnitude of phenotypic selection on reproductive and worker traits is required. Interestingly, some worker-associated genes in our dataset showed evidence of positive selection (Figs. S10, S11; Table S8), and α, the estimated proportion of substitutions fixed by positive selection for worker-associated genes, was greater than zero (Fig. 1C; bootstrap p < 0.001) and similar to the genomic background rate for non-differentially expressed genes (Fig. 1C; bootstrap p = 0.92). Some worker traits may simply experience strong phenotypic selection (e.g., on the number or survival of new sibling queens), overcoming the dilution effect of kin selection (Linksvayer and Wade 2009; Linksvayer and Wade 2016). In some species, phenotypic selection may even act more strongly on worker traits, as suggested by a recent honey bee population genomic study that found evidence that a set of 90 worker-upregulated genes experienced stronger selection than 79 queen-associated genes (Harpur et al. 2014).

Our study identified thousands of genes that have putatively been shaped by kin selection, and hence reveals the promise of identifying genome-wide signatures of social evolution. Our study lends support to the notion that social traits may have distinct genetic and evolutionary features (Mikheyev and Linksvayer 2015), even though the evolution of complex social traits such as caste are mainly based on the recruitment of ancient genes. Our results thus help to tie together previous sociogenomic studies, which have been motivated by concepts from Evolutionary Developmental Biology (Toth and Robinson 2007) and have stressed the importance of either highly conserved (Toth and Robinson 2007; Woodard et al. 2011; O’Connell and Hofmann 2012; Berens et al. 2015) or novel genes (Johnson and Tsutsui 2011; Ferreira et al. 2013; Feldmeyer et al. 2014; Sumner 2014; Jasper et al. 2016) for social evolution, with population genetic models based on well-established social evolution theory (Hamilton 1964; Linksvayer and Wade 2009; Hall and Goodisman 2012; Linksvayer and Wade 2016).

## METHODS

### Study design and sampling procedure

In order to collect a time series of developing worker and reproductive larvae, and also adult workers and queens, we set up a sacrifice study in which 30 total replicate experimental colonies were assigned to either a queen present or queen absent treatment and then sampled at one of five time points corresponding to larval developmental stages. Queen removal stimulates the production of new reproductives (i.e. new queens and males) (Edwards 1987; Schmidt et al. 2010) so that following queen removal, a portion of young brood (eggs and 1st instar larvae) are reared as reproductives, whereas all older brood are reared as workers. We also randomly assigned each experimental colony to one of five time points (L1-L5), corresponding to five larval developmental stages.

The timing of sampling for colonies in both treatments was based on the current age of the youngest larvae present in the queen removed treatment colonies, which corresponded to brood that were eggs at the time of queen removal. Thus, we sampled the first set of colonies assigned to stage L1 approximately five days after creation, at which point nearly all eggs in queen removed colonies had hatched into 1^st^ instar larvae. Colonies assigned to subsequent stages (L2-L5) were sampled in intervals of 3-4 days, yielding samples of colonies with L2, L3, L4, and L5 larvae. We collected the following samples from each colony: for queen present colonies, we collected worker larvae, adult worker foragers, and adult worker nurses; for queen absent colonies, we collected both worker and reproductive larvae, adult worker foragers, and adult worker nurses observed feeding worker larvae as well as adult worker nurses observed feeding reproductive larvae.

Ten individuals of each sample type were collected and pooled into a single sample. Each individual was immediately flash-frozen in liquid nitrogen after collection. Adult worker heads and gasters (i.e. the last four abdominal segments) were collected separately and removed from the body while frozen. To collect adult queen head and gaster samples, 10 mature egg-laying queens approximately 4 months old were collected from three of the genetically homogeneous stock colonies used to create the experimental colonies and processed in the same manner as adult worker samples.

### RNA sequencing and mapping

We extracted RNA using RNeasy kits in accordance with manufacturer’s instructions. 25 samples were removed due to contamination or degradation, as detected by an Agilent 2100 Bioanalyzer or poor yield (<50 ng RNA). After excluding these samples, we prepared 161 cDNA sequencing libraries using poly-T capture of messenger RNA and subsequent full-length amplification, as in Aird *et al.* (2013). For quality control and to estimate the dynamic range of the sequencing experiment, we added two ERCC92 (Thermo Fisher Scientific Inc.) spike-in mixes to total RNA, with half the samples randomly receiving one or the other mix. Sequencing of the cDNA libraries was performed on an Illumina HiSeq 2000 in SE50 mode at the Okinawa Institute of Science and Technology Sequencing Center. Reads were mapped to the assembly and NCBI version 2.0 gene models (Mikheyev and Linksvayer 2015) using RSEM (Li and Dewey 2011) to obtain expected counts and fragments per kilobase mapped (FPKM).

### Differential expression analysis

We removed genes with FPKM < 1 in at least half the samples of all three tissues (head, gaster, and larvae) from further analysis. We removed two samples from further analysis due to suspected contamination (see Supplemental Table 1 for numbers of samples used for subsequent analysis). We performed differential expression analysis using edgeR (Robinson and Oshlack 2010) with a GLM-like fit to the count data (McCarthy et al. 2012). In order to identify worker-upregulated and reproductive-upregulated genes, we performed differential expression analysis separately by larval stage and adult sample type, across all larval stages, and across all larval stages and adult samples together. We performed subsequent analyses using the sets of genes that had an overall average effect of caste across all larval and adult samples. We assumed that these genes were most tightly associated with worker versus reproductive function, and hence shaped primarily by indirect (i.e. kin) selection versus direct selection.

### Population genomic analysis

We constructed genomic sequencing libraries for 22 single-worker specimens of *M. pharaonis* and one outgroup worker sample of *Monomorium chinense*. We chose the ingroup samples to maximize geographic coverage and to provide a representative sample of standing genetic diversity in this species. Sequencing libraries were made using Illumina Nextera kits and sequenced on an Illumina HiSeq 2000 instrument. *M. pharaonis* reads were mapped to the reference using bowtie 2 in very sensitive local mode (Langmead and Salzberg 2012), while the *M. chinense* samples were mapped using NextGenMap (Sedlazeck et al. 2013), which offers more sensitivity for divergent sequences. Subsequently, variants were called separately using GATK, FreeBayes and Samtools (Li et al. 2009; McKenna et al. 2010; Garrison and Marth 2012). These variant call sets were converted to allelic primitives using GATK, and combined into a high credibility set using BAYSIC (Cantarel et al. 2014). We subsequently removed indels, any sites with more than two alleles, with more than 10% missing data, and any with a site quality lower than a phred score of 40 to produce the final variant call set. The effect of each variant (synonymous vs. nonsynonymous) was determined using SnpEff (Cingolani et al. 2012). We then used the resulting table of numbers of synonymous polymorphisms (*P_S_*) and substitutions (*D_S_*) and nonsynonymous polymorphisms (*P_N_*) and substitutions (*D_N_*) for input for McDonald-Kreitman (McDonald and Kreitman 1991) test-based software for estimating population genetic parameters and inferring signatures of selection (Welch 2006; Eilertson et al. 2012).

The McDonald-Kreitman test can be extended to estimate α, the proportion of amino acid substitutions that are fixed by positive selection (Bierne and Eyre-Walker 2004)(Smith and Eyre-Walker 2002; Welch 2006), as a powerful way to study genome-wide rates of adaptive molecular evolution. We estimated α for worker-upregulated, reproductive-upregulated, and non-differentially expressed genes. We used a maximum likelihood estimator developed in the software package MKtest2.0 (Welch 2006; Obbard et al. 2009) (available at http://sitka.gen.cam.ac.uk/research/welch/GroupPage/Software.html, last accessed 1 July, 2016).

### Comparative genomic phylostratigraphy analysis

We constructed phylostratigraphic maps for *M. pharaonis*, as well as two species with higher quality genomes (*Apis mellifera*, and *Drosophila melanogaster*), following previously developed methods (Domazet-Loso et al. 2007; Domazet-Lošo and Tautz 2010; Quint et al. 2012; Drost et al. 2015). Phylostrata were defined for each species according to the NCBI taxonomy database (Table S4; Fig. S6). We constructed a target database by adding recently sequenced hymenopteran genomes to a database that was recently used in a phylostratigraphy study of animal and plant development (Drost et al. 2015). Species-specific amino acid sequences were downloaded from RefSeq (Pruitt et al. 2012), last accessed 11 July, 2016. Each amino acid sequence at least 30 amino acids long for each of the three species were used as a query against the target database using BLASTp (version 2.2.25). Transcripts were assigned to the oldest phylostrata containing at least one BLAST hit with an E-value below 10^−5^ for the given transcript. If no BLAST hit with an E-value below 10^−5^ was found, the transcript was placed in the youngest, species-specific phylostrata (e.g., *Monomorium pharaonis*). Genes were assigned to phylostrata based on the phylostrata of their longest transcript isoform. To verify that our results did not depend on the E-value threshold we used, we also constructed a map for *M. pharaonis* using a very liberal E-value threshold of 10^−1^ as well as the default, much more conservative 10^−5^ threshold (Quint et al. 2012).

To compare mean phylostrata between queen- and worker-associated genes, we used generalized linear models with poisson residuals (or quasipoisson for overdispersed models). We used both raw phylostrata (after removing any phylostrata with zero genes; PS in Table S4), as well as phylostrata condensed into 6 main categories because many categories had few genes (Fig. S6): cellular organisms, eukaryotes, bilaterian animals, insects, hymenopterans, and ants (“Condensed PS1” Table S4). To compare the relative contribution of phylostrata to worker- and reproductive-associated genes, we constructed contingency tables, used omnibus Chi-square tests, calculated standardized Pearson’s residuals to explore the contribution of each cell to the omnibus test, and presented the results using mosaic plots with the “vcd” R package (Friendly and Meyer 2015) and the 6 Condensed PS1 categories. The significance of enrichment for individual cells was assessed with standardized Pearson residuals, where residuals with an absolute value > 2 have an approximate p-value < 0.05, and residuals with an absolute value > 4 have an approximate p-value < 0.001 (Friendly 1994)(Friendly and Meyer 2015).

### Gene Ontology enrichment analysis

We calculated GO term enrichment of categories of identified worker-associated and reproductive-associated genes, as well as worker-associated and reproductive-associated genes grouped by phylostrata, using the R package “GOstats”, with a cut-off p-value of 0.05 (Falcon and Gentleman 2007).

### Statistical analyses

All statistical analyses and figures were made with R version 3.1.2.

### Data deposition

Raw sequencing reads will be deposited in DDBJ bioproject PRJDB3164. Count and FPKM data, and a .csv file summarizing all analyses for each locus are available as Supplementary Materials.

## ACKNOWLEDGEMENTS

This work was supported be supported by the National Science Foundation (grant number IOS-1452520 to T.A.L.) and Okinawa Institute for Science and Technology subsidy funding to A.S.M..

## Supplementary Material

### Supplemental Methods

#### Study species

*Monomorium pharaonis* has the following suite of traits that make it suitable for the current study: unlike most ants and other hymenopteran social insects, which have facultatively sterile workers that can lay male-destined eggs under some conditions, *Monomorium* workers are obligately sterile (Hölldobler and Wilson 1990), so that genes exclusively expressed by *M. pharaonis* workers can only have indirect fitness effects; *M. pharaonis* colonies are readily experimentally induced to shift from producing only new workers, to producing a mixture of new workers and reproductives (i.e. queens and males), by removing current egg-laying queens (Schmidt et al. 2010)(Edwards 1987); worker- and reproductive-destined larvae can be morphologically distinguished at an early developmental stage (Fig. 1A) (Berndt and Kremer 1986a); controlled crosses can readily be made in the lab and hundreds of colonies kept across generations; and aggression between workers from different colonies is transient so that the genetic makeup of colonies can be experimentally controlled.

#### Study design and colony setup

The study was run in three total blocks, each separated by three weeks, starting in April 2014. For each block, we did the following:

1. We created a genetically homogeneous source by mixing at least 10 large stock colonies, which themselves had been repeatedly mixed across generations (note that unlike most ants, *M. pharaonis* colonies display at most transient aggression following colony mixing).
2. From this source, we allocated 0.5 mL of mixed brood and workers to each replicate experimental colony, resulting in colonies with ∼300-400 workers and ∼300-400 brood of various stages (i.e. eggs, larvae, pupae).
3. We randomly assigned half of the experimental colonies to a queen present treatment, where queen number was standardized to 10 queens, and the other half to a queen absent treatment, where all queens were removed. Queen removal stimulates the production of new reproductives (i.e. new queens and males) (Schmidt et al. 2010)(Edwards 1987) so that following queen removal, a portion of young brood (eggs and 1st instar larvae) are reared as reproductives, whereas all older brood are reared as workers.
4. We also randomly assigned each experimental colony to one of five time points (L1-L5), corresponding to five larval developmental stages (see below).

All colonies were maintained at 27 ± 1 °C and 50% humidity, and fed twice weekly with dried mealworms (*Tenebrio molitor*) and an agar-based synthetic diet (Dussutour and Simpson 2008). Experimental colonies were maintained in glass nests made of two pieces of 4 cm × 6 cm glass separated by 1.5 mm strips of plastic. All surveys and sampling were performed using dissecting microscopes.

#### Sampling procedure

Note that adult *M. pharaonis* workers performing foraging or nursing tasks have been shown to have divergent whole-body gene expression profiles (Mikheyev and Linksvayer 2015), and we collected forager and nurse samples across social contexts (queen presence and larval stage) in order to be able to confidently identify genes that are upregulated in worker tissues.

On the day designated for sample collection, colonies were placed in petri dishes, surveyed, and then lightly anesthetized using carbon dioxide to prepare the colony for sample collection. While anesthetized, the top glass pane of the nest was removed to enable collection of nursing workers, the petri dish was covered with a lid to minimize disturbance from air flow, and the petri dish was placed on a heating pad, kept at 27 °C, to maintain a constant temperature throughout sample collection. Colonies were left undisturbed for 30 minutes to recover from anesthesia prior to sample collection. Foragers were collected when observed collecting food outside the nest, and nurses were collected when observed nursing the appropriate larval stage. For example, for the L2 sample, worker nurses were collected when witnessed feeding a 2^nd^ instar worker larva, and reproductive nurses when witnessed feeding a 2^nd^ instar reproductive larva. After all foragers and nurses were collected, colonies were lightly anesthetized again and larvae of the appropriate stage were collected.

Larval instars were determined by overall size, shape, and especially hair presence and morphology (Berndt and Kremer 1986b). *M. pharaonis* larvae have three distinct larval instars (Berndt and Kremer 1986b), but because the vast majority of growth occurs in the third larval instar, we divided the third instar into three separate stages (L3-L5) based on size (Fig. 1A). Third instar worker larvae were defined as the L3 stage until they reached 0.75x the length of a worker pupa, L4 stage up to 1x the length of worker pupae, and L5 thereafter. Reproductive larvae are hairless (Berndt and Kremer 1986b) and can be differentiated from worker larvae starting at the 2^nd^ larval instar. Reproductive larval stages were defined as follows: 2^nd^ instar (i.e. L2) until 0.5x the length of a worker pupa, L3 from 0.5-1x the length of a worker pupae, L4 from 1-1.5x the length of a worker pupa, and L5 thereafter.

Note that it is not possible to morphologically distinguish between male and queen larvae, so that the reproductive larvae we collected included both males and queens. *M. pharaonis* colonies produce female-biased reproductive sex ratios (e.g., 0.739 female [queen/(queen+male)], interquartile range = 0.028, based on 39 colonies, (Schmidt et al. 2010)), and furthermore, newly eclosed adult queens are on average 1.42 times as large as males (1.359 mg wet mass versus 0.955 mg, based on samples of 247 newly eclosed queens and 235 males, (Schmidt et al. 2010)). Thus, we estimate that approximately 80% of the sampled larval tissue came from queens, so that the transcriptomic profiles we observed for our reproductive larvae samples mostly reflected queen larvae.

We separately collected adult head and abdominal tissues to increase the likelihood of detecting differentially expressed genes associated with adult behavior (e.g., genes upregulated in worker brains) and function (e.g., genes upregulated in queen reproductive tissues).

#### Differential expression analysis

For stage-specific analyses, caste was the only factor included. For analyses including all larval samples or all adult and larval samples, we used a model with caste and stage as fixed factors. For comparisons only considering larval stages, we included batch as an additional factor, but we could not include batch in comparisons including adult samples because the queen samples were not collected within the same blocked design.

#### Population genomic analyses

The McDonald-Kreitman test (McDonald and Kreitman 1991) uses both polymorphism and substitution data (e.g., an excess of nonsynonymous substitutions relative to polymorphisms; *D_N_*/*D_S_* > *P_N_*/*P_S_*) to infer the fixation of advantageous mutations by positive selection (Bierne and Eyre-Walker 2004). This approach is more powerful at identifying signatures of positive selection than using only substitution data for divergent lineages (i.e. with *d_N_*/*d_S_*, which is *D_N_*/*D_S_* weighted by the total numbers of nonsynoymous and synonymous sites), because elevated *d_N_*/*d_S_* estimates can arise either from positive selection or relaxed purifying selection.

MKtest2.0 was especially suitable for our needs because it can simultaneously estimate α for different gene categories (Obbard et al. 2009), instead of only providing a single genome-wide estimate. Besides estimating α, MKtest2.0 can estimate other population genetic parameters, including selective constraint, described by 1-*f*, the proportion of non-synonymous mutations that experience strong purifying selection, as well as neutral diversity, and neutral divergence. Because we were interested in comparing patterns of selection experienced by worker- and reproductive-upregulated genes, and to avoid overparameterized models, we focused on α and *f* and kept other parameters at default values (i.e. a single global value). We compared model estimates for α and *f* and model fit statistics using a single genome-wide estimate (i.e. the default), separate estimates for each of three categories (i.e. reproductive-associated, worker-associated, and NDE), and for *f*, we also considered separate estimates for each locus. Because models with gene-specific *f* estimates were best (Table S6), we focus on the estimates from these models in the main text, although we observed similar patterns for other parameters as well. We estimated 95% confidence intervals for α with the bootstrapping feature in MKtest2.0 (i.e. as 95% bootstrap intervals around the mean, based on 1,000 bootstrap replicates across genes). We also used bootstrapping to determine p-values for the hypothesis that reproductive-associated genes have α greater than worker-associated genes. Similarly, we used bootstrapping to determine p-values for the hypothesis that caste-associated genes grouped by phylostrata had α greater than zero. We used beta regression (R package “betareg”; (Cribari-Neto and Zeileis 2010)), to compare mean *f* estimates between worker-associated and queen-associate genes.

In addition to estimating α across groups of genes in order to compare rates of adaptive molecular evolution at these genes, it is possible to estimate selection coefficients separately for each locus and then compare mean estimated selection coefficients. The software SnIPRE (Eilertson et al. 2012), Selection Inference using Poisson Random Effects, is a Bayesian implementation of the McDonald-Kreitman test and seeks to estimate several population genetic parameters similar to those estimated by MKtest2.0, including the selection coefficient acting on every gene weighted by effective population size (γ = 2*N_e_s*). We used SnIPRE to estimate γ for each gene and compared mean γ for the categories of caste-associated genes we identified. Specifically, we used a generalized linear model (glm) to compare mean BSnIPRE.est, a normalized estimate of γ produced by SnIPRE, for worker-associated, reproductive-associated, and NDE genes.

We also used SnIPRE to categorize genes as experiencing positive, neutral, or negative selection. We also classified genes as experiencing selection using the standard McDonald-Kreitman test (McDonald and Kreitman 1991), by plotting log(p-value) from this test versus the -log(*NI*) (Li et al. 2008), where NI is the neutrality index. We used an unbiased estimator for NI (Stoletzki and Eyre-Walker 2010). Genes above a threshold p-value from the McDonald-Kreitman test with a positive -log(*NI*) were categorized as having experienced positive selection, and those with a negative -log(*NI*) were categorized as having experienced negative selection https://paperpile.com/c/JsHRHS/PlAhk (Li et al. 2008). We used both a liberal cutoff of 0.05 for the nominal p-value from the McDonald-Kreitman test, and also a much more conservative Bonferroni-corrected p-value cutoff (Li et al. 2008) based on the 5,674 genes included in the analysis (genes with zeros for any of the four counts *D_N_*, *D_S_*, *P_N_*, *P_S_* were excluded because such zeros lead to undefined *NI*).

#### Comparative genomic phylostratigraphy analysis

Phylostratigraphy attempts to estimate the evolutionary age of protein-coding genes in a focal species by identifying the distribution of their homologs across the tree of life (Domazet-Loso et al. 2007; Domazet-Lošo and Tautz 2010; Quint et al. 2012; Drost et al. 2015). The approach sorts genes into hierarchical phylostrata (PS), based on the oldest BLAST hit of their amino acid sequence. For example, a gene from an ant species with identified orthologs across all eukaryotes is assumed to be much older than a gene with only identifiable orthologs in other closely related ant species. We were interested in estimating the relative ages of the the sets of worker-upregulated, reproductive-upregulated, and non-differentially expressed genes that we identified, and were less interested in the precise age estimates. Indeed, even though phylostratigraphy has been widely used (Domazet-Loso et al. 2007; Domazet-Lošo and Tautz 2010; Quint et al. 2012; Drost et al. 2015), the precise age estimates can be influenced by several factors, including parameters used to define homology in BLAST (e.g., threshold gene length, threshold E-value, database size, etc.) (Moyers and Zhang 2016)(Moyers and Zhang 2015). Furthermore, because homologous sequences in BLAST are generally required to span relatively small lengths (i.e. 30 amino acids) (Quint et al. 2012; Drost et al. 2015), small portions of a gene can impact the phylostrata assigned.

#### Statistical analyses and figures

All statistical analyses and figures were made with R version 3.1.2, using packages “ggplot2”, “gplots”, “scales”, “stats”, “plyr”, “ggdendro”, “gridExtra”, “tidyr”, “plyr”, “vcd”, “vcdExtra”, “Vennerable”, “data.table”, “edgeR”, “myTAI”, “betareg”, and “GOstats”. Complete R scripts used in the analyses will be included in the final publication. For Figure 1A, we collected representative worker and reproductive larvae from each stage (Berndt and Kremer 1986b) and arranged them in a series in order to produce a figure representing the developmental time series we used.

**Fig. S1.**
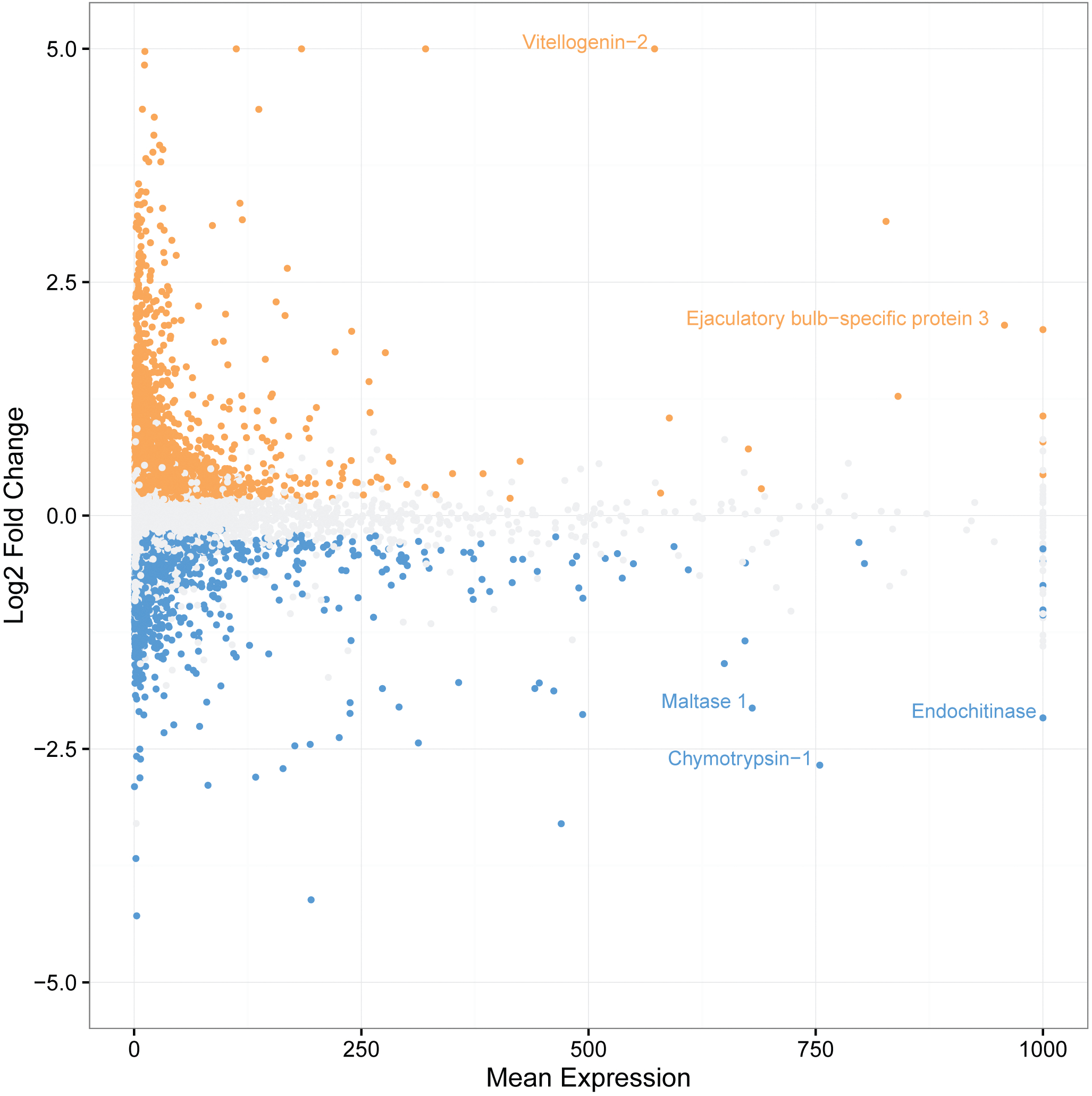
Log2 fold change (Reproductive/Worker) as a function of the mean of worker and reproductive expression (FPKM) across all larval stages and adult samples. Genes with annotation information and FPKM > 500 and a LogFC of a greater magnitude than 2.5 are labeled. Genes that are colored are differentially expressed, with a main effect of caste across samples: orange = reproductive-upregulated; blue = worker-upregulated; grey = nondifferentially expressed (NDE). For plotting purposes, genes with a log2 fold greater than (less than) 5 (-5) assigned a value of 5 (-5). Genes with greater mean expression than 1000 FPKM assigned a value of 1000 FPKM.

**Fig S2.**
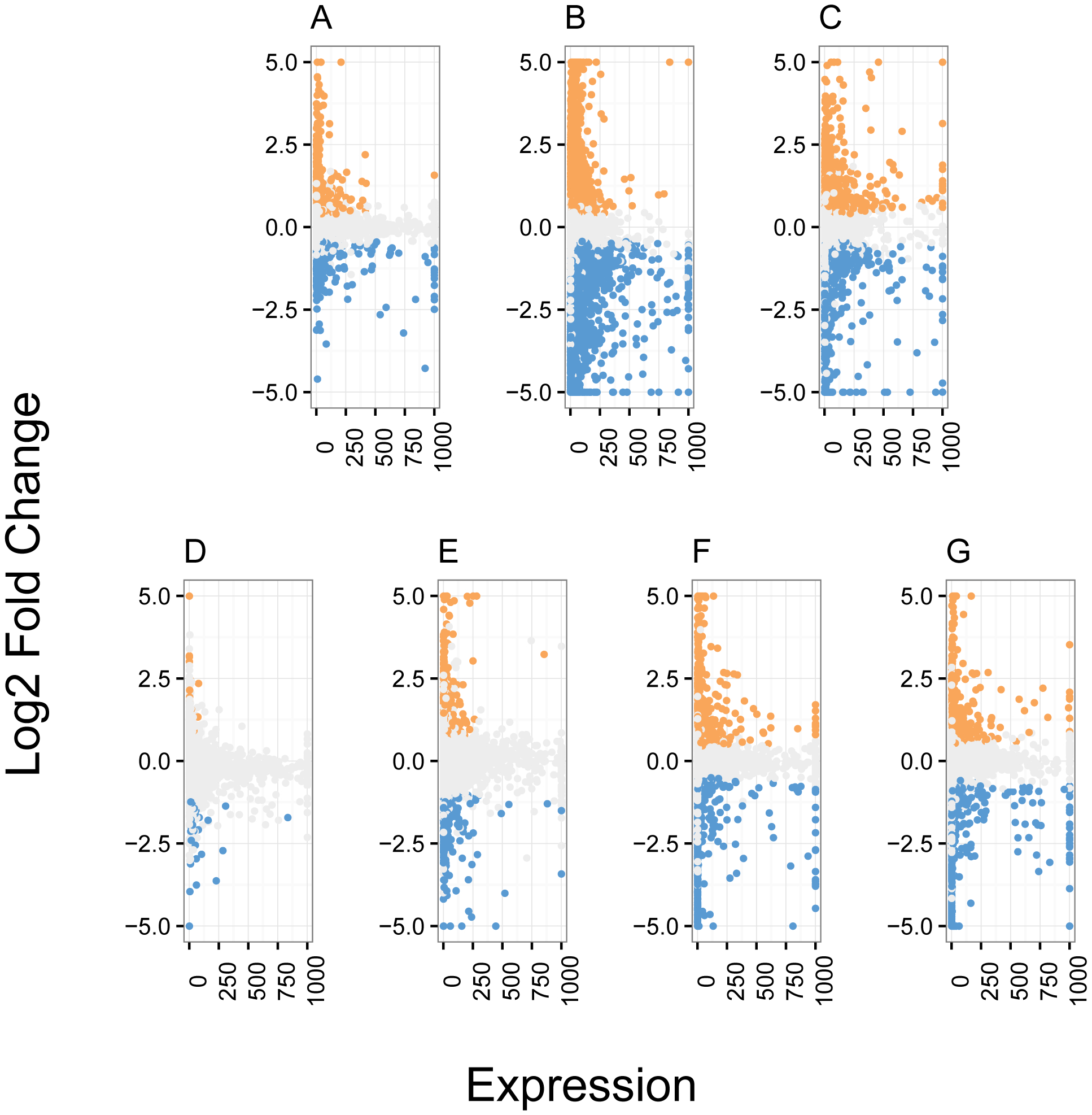
Log2 fold change (Reproductive/Worker) as a function of the mean of worker and reproductive expression (FPKM) for specific larval stages or adult samples. Results are from (**A**) across larval stages L2-L5, (**B**) adult head, (**C**) adult gaster, (**D**) L2, (**E**) L3, (**F**) L4, and (**G**) L5. Genes that are colored are differentially expressed, with a main effect of caste across samples: orange = reproductive-upregulated; blue = worker-upregulated; grey = NDE. For plotting purposes, genes with a log2 fold greater than (less than) 5 (-5) were assigned a value of 5 ( -5). Genes with greater mean expression than 1000 FPKM were assigned a value of 1000 FPKM.

**Fig. S3.**
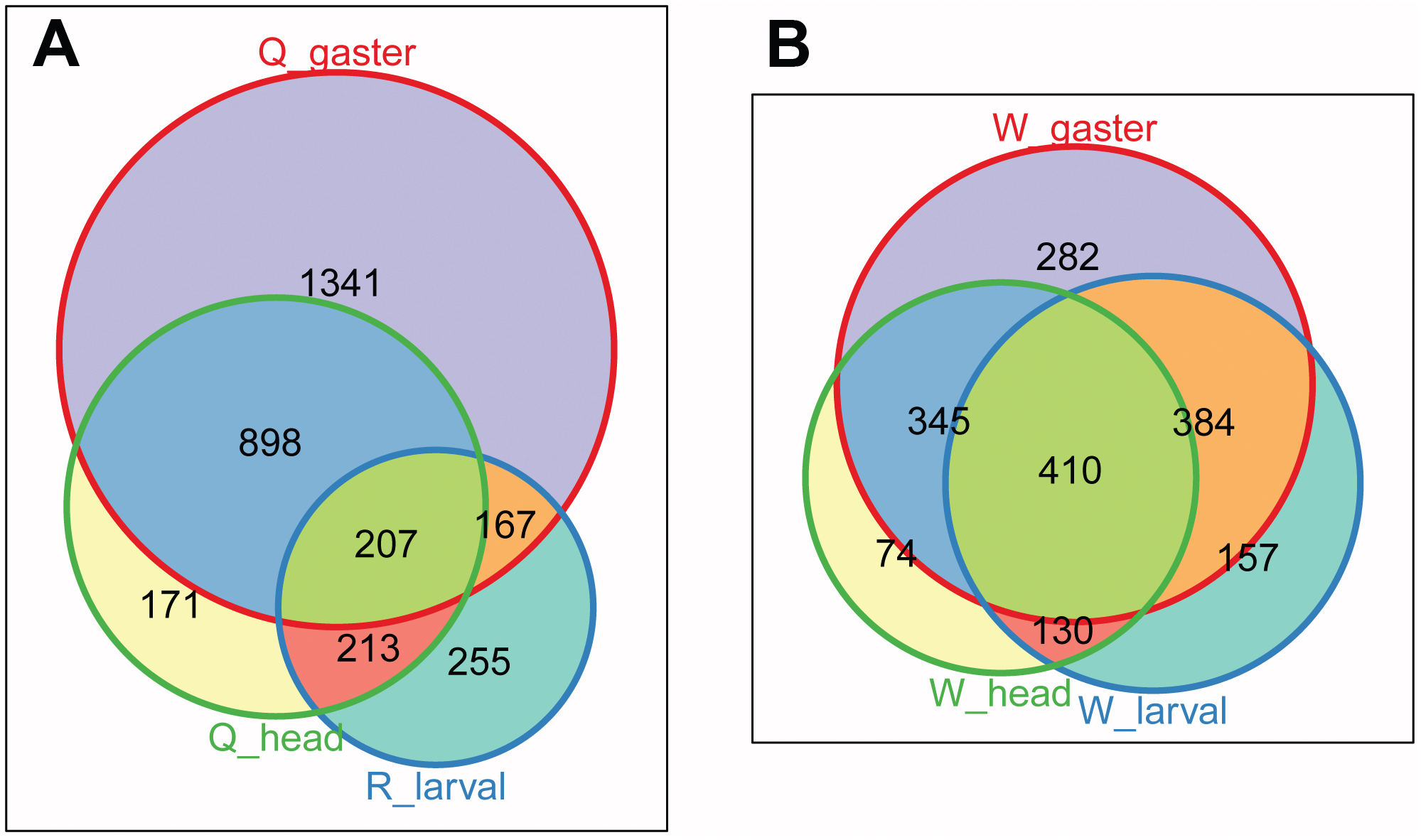
Weighted three-set Venn diagrams showing the contribution of: **A.** queen gaster-upregulated genes, queen head-upregulated genes, and the union of all reproductive larvae-upregulated genes (for L2-L5) to the set of reproductive-upregulated genes with a main effect across all samples; **B.** Worker gaster-upregulated genes, worker head-upregulated genes, and the union of all worker larvae-upregulated genes (for L2-L5) to the set of worker-upregulated genes with a main effect across all samples. Note that the set of reproductive-upregulated genes is dominated by genes upregulated in adult queen tissues, with 80% of reproductive-upregulated genes upregulated in queen abdominal (i.e. gaster) tissue, and 46% in queen head tissue. 41% (i.e. 1341/3252) are only upregulated in queen gasters, and not in any other tissue. In contrast, the set of worker-upregulated genes is more evenly composed of genes upregulated in worker gaster, head, and larval samples.

**Fig. S4.**
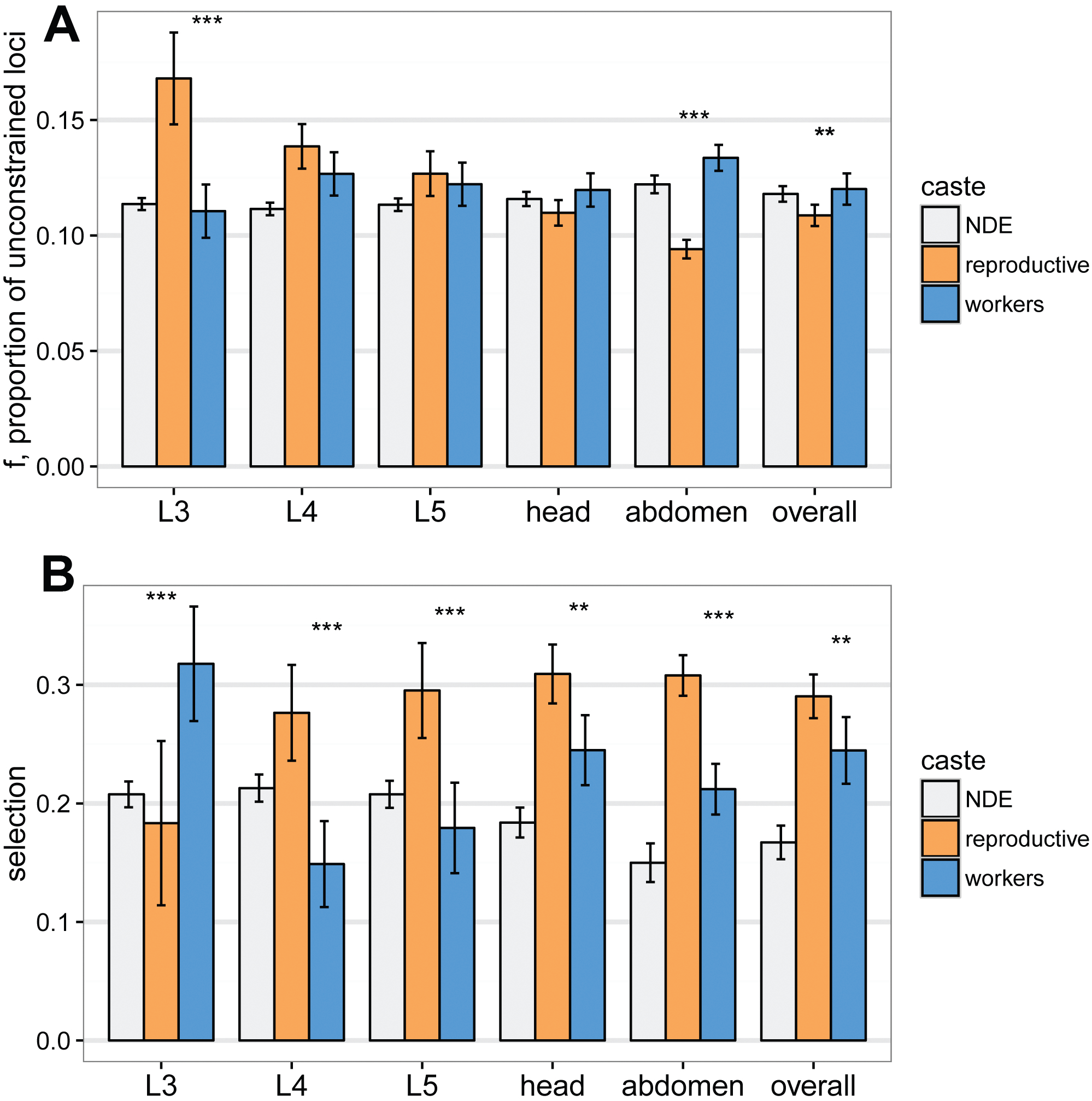
Per-locus selective constraint and selection coefficients across samples. (**A**) Reproductive-upregulated genes with a main effect across all samples (“overall”) and reproductive-upregulated genes from queen abdomens had higher mean selective constraint (=lower *f*) than worker-upregulated genes. Locus-specific *f* estimates were made with MKtest2.0. (**B**). Except for the L3 comparison, reproductive-upregulated genes in all comparisons have higher mean selection coefficients estimated by SnIPRE (glm using the normalized estimate BSnIPRE.est). For L3, worker-associated genes have a higher estimate than reproductive associated genes. * p < 0.05, ** p < 0.01, *** p < 0.001.

**Fig. S5.**
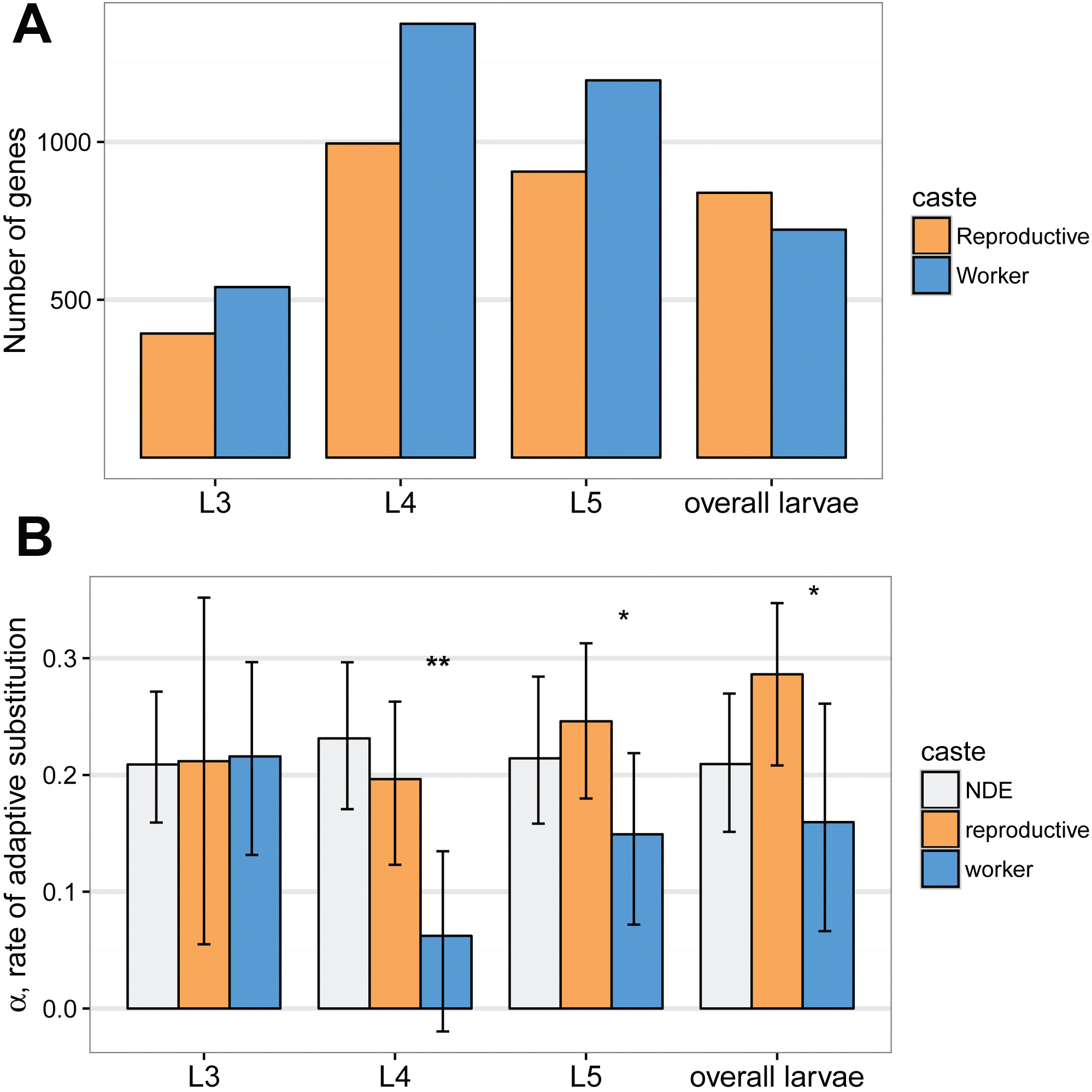
(**A**) Number of differentially expressed genes for only larval samples (L3-L5) as well as “overall larvae”, genes with a main effect of caste across larval samples. (**B**) Reproductive-associated genes have higher α, the proportion of amino acid substitutions fixed by positive selection, for larval genes, except at the L3 stage. * p < 0.05, ** p < 0.01.

**Fig. S6.**
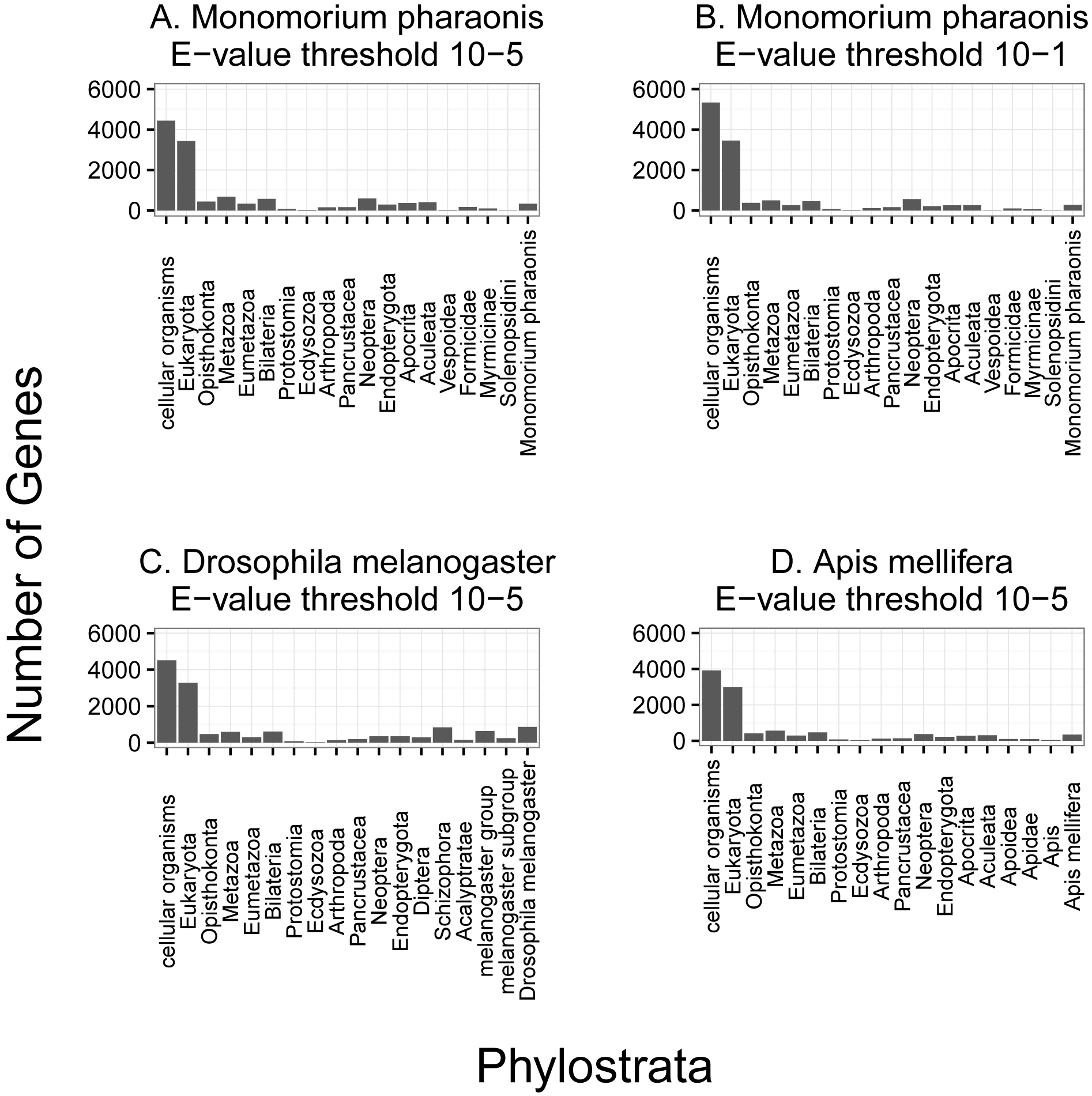
Number of genes in each phylostrata, as defined by the NCBI taxonomy database for: (**A**) *M. pharaonis* (E-value = 1 × 10^−5^), (**B**) *M. pharaonis* (E-value = 1 × 10^−1^), (**C**) *D. melanogaster* (E-value = 1 × 10^−5^), (**D**) *A. mellifera* (E-value = 1 × 10^−5^). The overall distribution is similar for all three species, with the majority of genes being ancient, and the pattern observed for *M. pharaonis* is consistent even when a very liberal threshold (E-value = 1 × 10^−1^) is used. BLASTp hits are made against a database containing nearly all species with curated genome annotations, with a minimum match length of 30 amino acids and a maximum E-value as listed above.

**Fig. S7.**
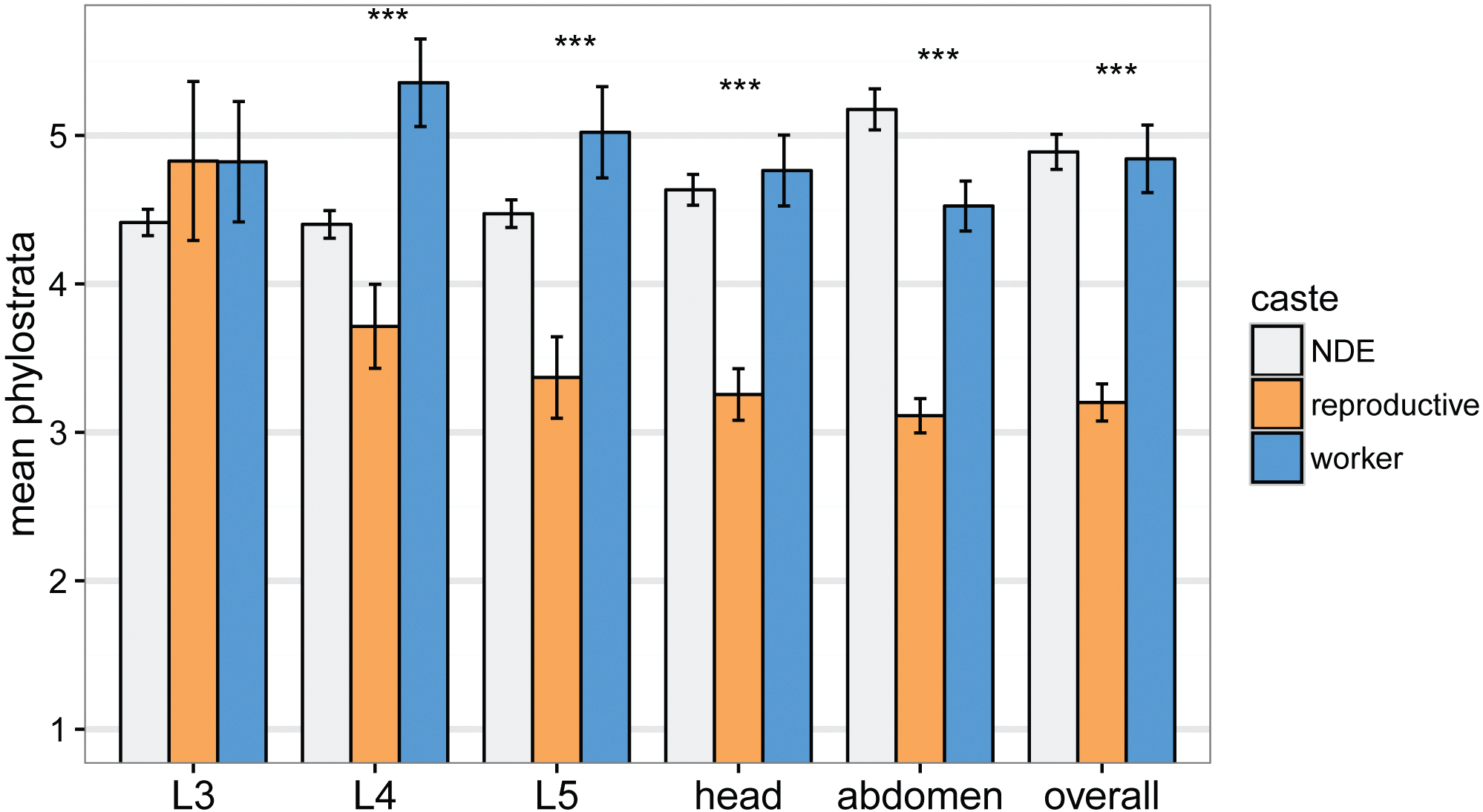
Using the original 19 phylostrata, reproductive-upregulated genes were older on average (i.e. lower mean phylostrata) for all comparisons except L3, the same result as when using grouped phylostrata (Fig. 1D). *** p<0.001.

**Fig. S8.**
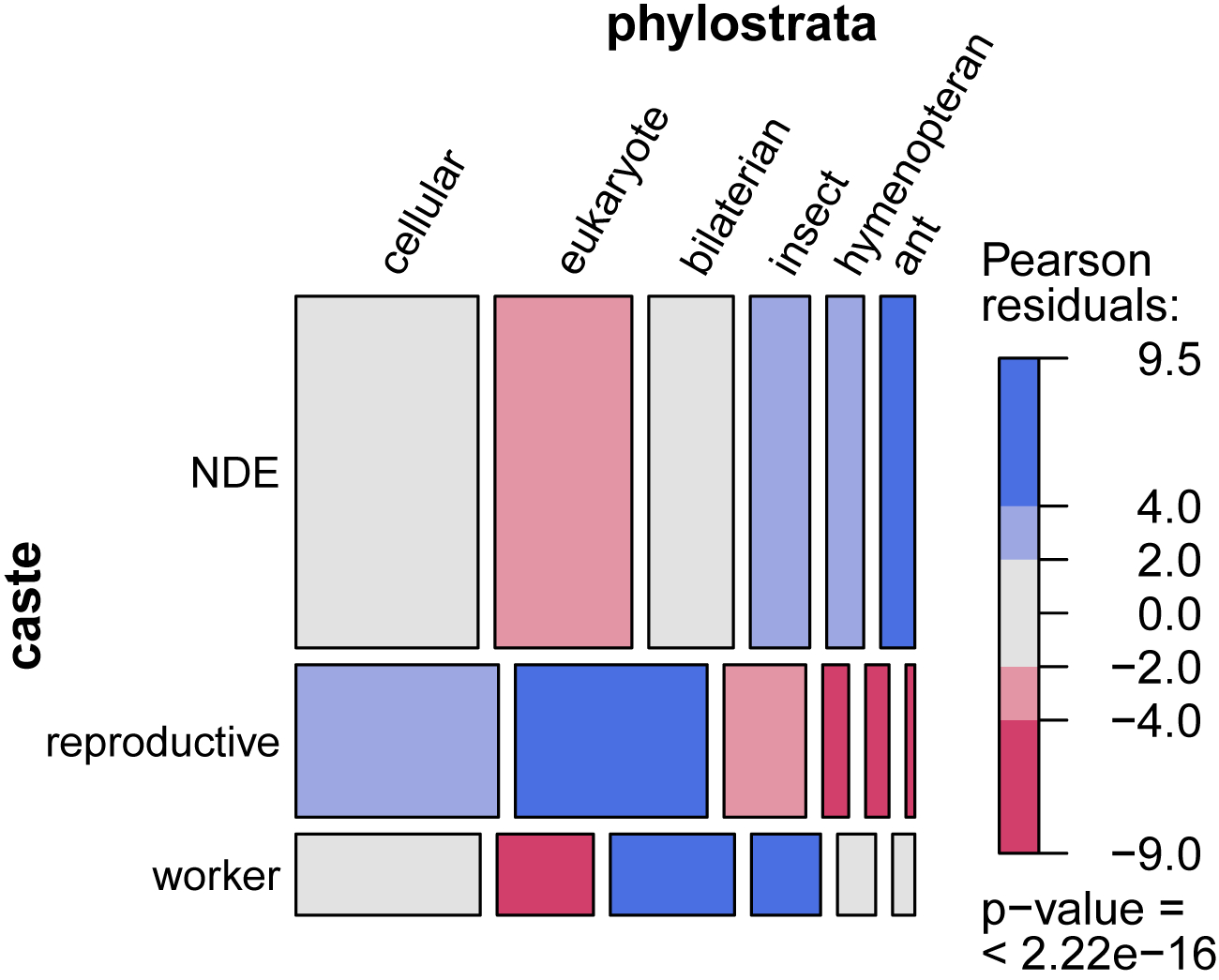
Mosaic plot showing the relative contribution of phylostrata to sets of reproductive-associated, worker-associated, and NDE genes. As when only considering worker- and reproductive-associated genes (Fig. 2C), reproductive-associated genes are enriched for the eukaryote phylostratum, but also for the cellular organism phylostratum. Similarly, worker-associated genes are enriched for bilaterian animal and insect phylostrata, but relative to NDE genes are no longer significantly enriched for the youngest two phylostrata (hymenopteran and ant) (Fig. 2C). The area of each cell is proportional to the number of genes in each caste and phylostrata category. Blue shading indicates overrepresentation (light blue p < 0.05, dark blue p < 0.001), and red-shading indicates underrepresentation (light red p < 0.05, dark red, p < 0.001), based on cell standardized pearson residuals.

**Fig. S9.**
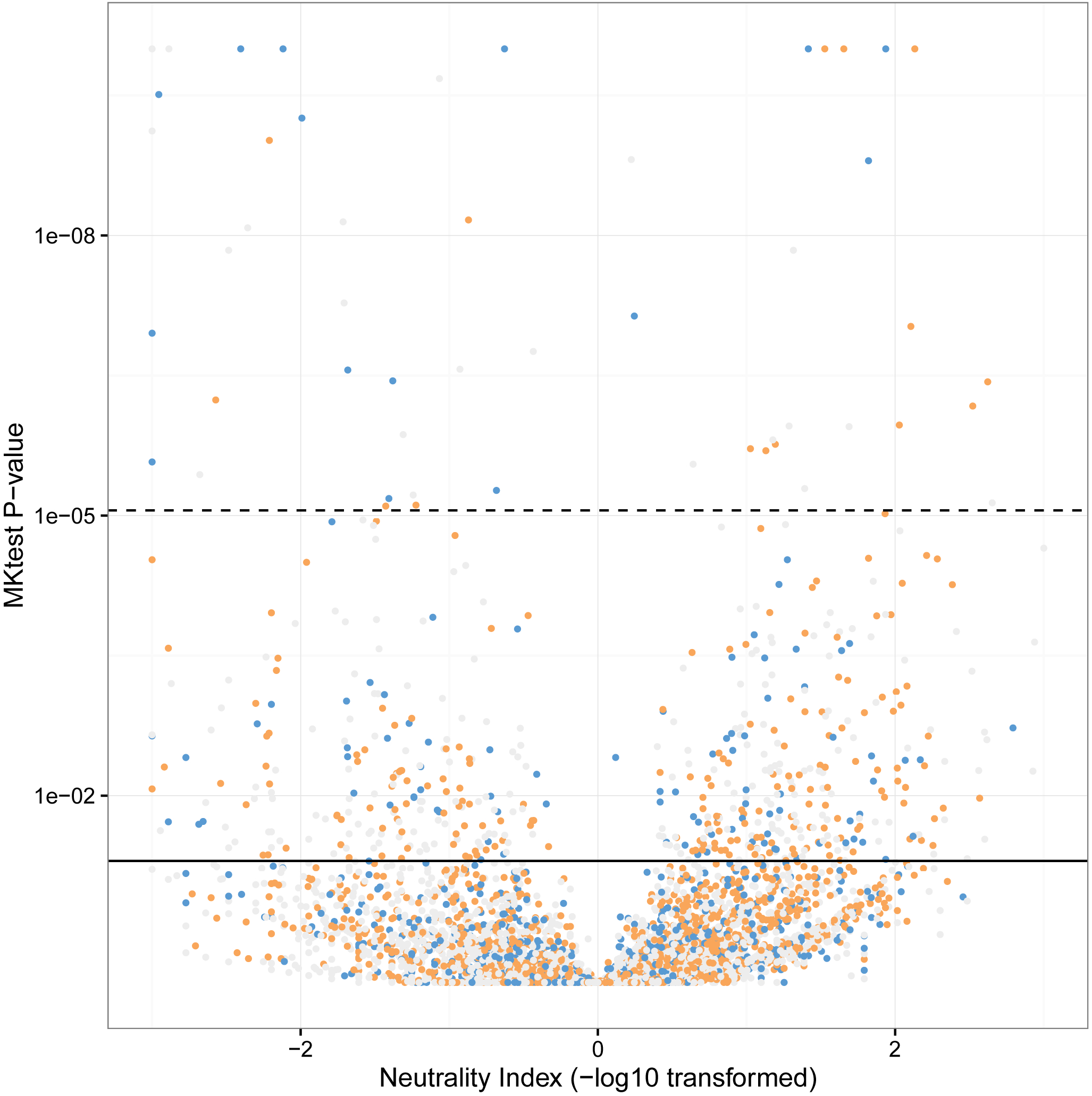
P-value as calculated from the McDonald-Kreitman test plotted against the neutrality index, which has been -log transformed, such that positive values indicate positive selection and negative indicate purifying/balancing selection. The solid black line indicates the nominal p-value (0.05) while the dashed line indicates the p-value after Bonferroni correction (N = 5674). Genes are colored by differential expression: grey = non-differentially expressed; orange = reproductive; blue = worker. For plotting purposes, genes with p-values less than 1 × 10^−10^ were assigned a p-value of 1 × 10^−10^, and those with a -log transformed neutrality index of greater (less) than 3 (-3) were assigned a value of 3 (-3).

**Fig. S10.**
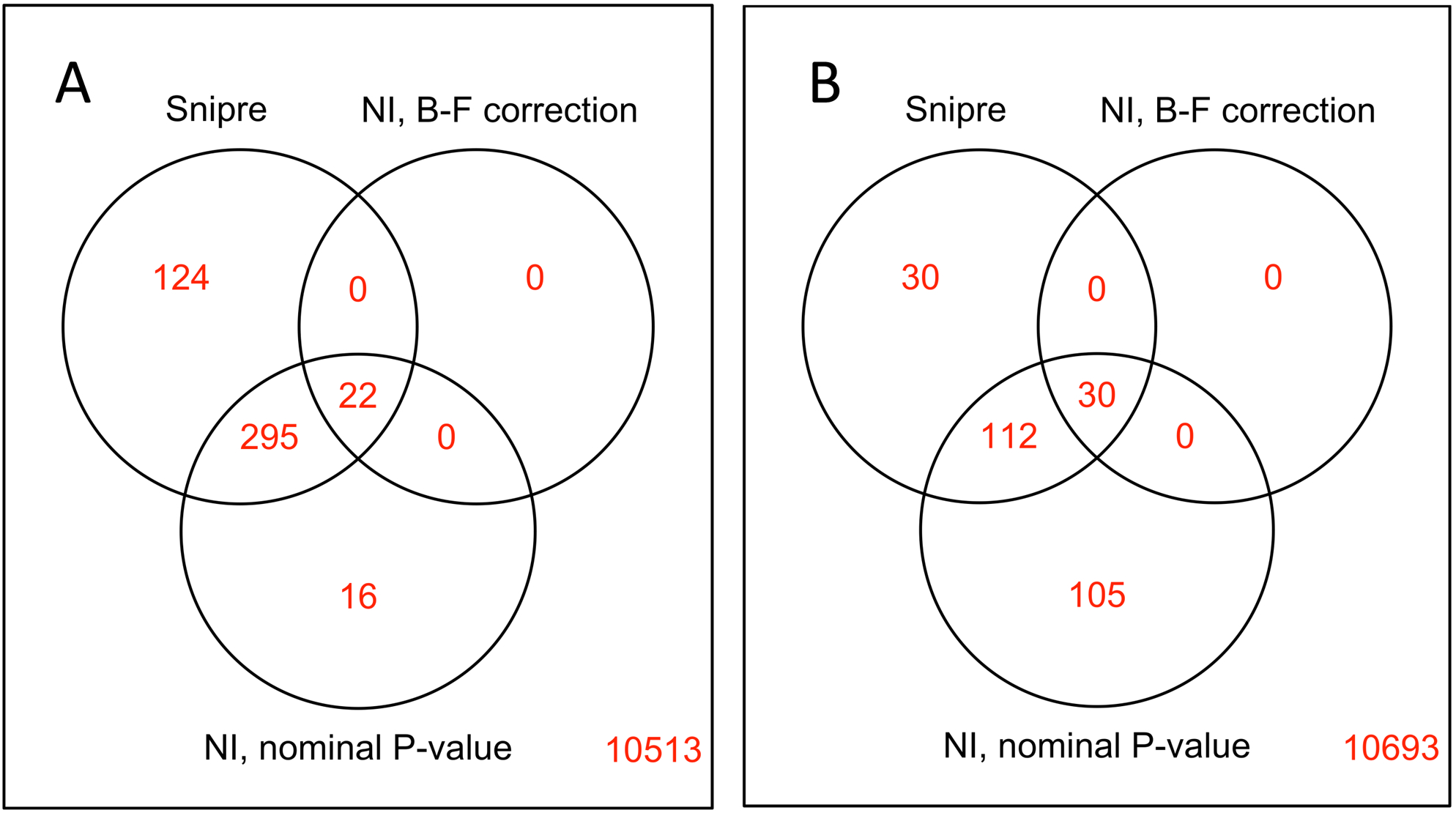
Overlap of A) positively and B) negatively selected genes as defined by SNiPRE and the Neutrality Index. Genes with negative values of -log10(Neutrality Index) and p-values less than 0.05 (for nominal) are defined by the “NI” method as under purifying selection, while such genes with positive -log10(Neutrality Index) values are assigned to the positive selection category. “NI, B-F correction” uses the same method but the p-value cutoff from the MKtest is adjusted for multiple comparisons using the Bonferroni procedure.

**Table S1.** Samples included in the study. Stages (L1 - L5) refer to the developmental stage of the larvae sampled at the particular stage of sample collection (see Sampling Procedure). Adult queen samples are marked “Other” because they were collected separately.

| **Sample** | **Queen Presence** | **L1** | **L2** | **L3** | **L4** | **L5** | **Other** | **Total** |
| --- | --- | --- | --- | --- | --- | --- | --- | --- |
| Forager Head | Present | 1 | 3 | 3 | 1 | 1 | 0 | **9** |
| Forager Gaster | Present | 2 | 2 | 3 | 2 | 2 | 0 | **11** |
| Forager Head | Absent | 2 | 3 | 3 | 3 | 2 | 0 | **13** |
| Forager Gaster | Absent | 2 | 3 | 2 | 3 | 2 | 0 | **12** |
| Worker Nurse Head | Present | 3 | 3 | 3 | 2 | 2 | 0 | **13** |
| Worker Nurse Gaster | Present | 3 | 3 | 3 | 3 | 2 | 0 | **14** |
| Worker Nurse Head | Absent | 2 | 3 | 2 | 2 | 3 | 0 | **12** |
| Worker Nurse Gaster | Absent | 3 | 2 | 3 | 2 | 3 | 0 | **13** |
| Reproductive Nurse Head | Absent | 0 | 2 | 2 | 2 | 3 | 0 | **9** |
| Reproductive Nurse Gaster | Absent | 0 | 3 | 3 | 3 | 3 | 0 | **12** |
| Worker Larva | Present | 0 | 3 | 3 | 3 | 3 | 0 | **12** |
| Worker Larva | Absent | 0 | 3 | 3 | 3 | 3 | 0 | **12** |
| Reproductive Larva | Absent | 0 | 3 | 3 | 3 | 3 | 0 | **12** |
| Adult Queen Head | Present | 0 | 0 | 0 | 0 | 0 | 3 | **3** |
| Adult Queen Gaster | Absent | 0 | 0 | 0 | 0 | 0 | 2 | **2** |

**Table S2.** Top 20 worker-upregulated genes, sorted by FDR. Differential expression calculated using glm-like model including caste and developmental stage as fixed effects. Negative values of Log2 fold change indicate higher expression in worker samples.

| Gene | Log2 Fold Change | P-Value | FDR | Description (SwissProt) |
| --- | --- | --- | --- | --- |
| **LOC105841041** | -1.83 | 2.62E-32 | 3.19E-30 | - |
| **LOC105837944** | -1.39 | 2.13E-26 | 1.67E-24 | - |
| **LOC105835450** | -0.83 | 4.82E-26 | 3.62E-24 | - |
| **LOC105835451** | -1.03 | 5.34E-23 | 2.96E-21 | Endothelin-converting enzyme 1 |
| **LOC105834581** | -1.09 | 1.82E-22 | 9.57E-21 | TPPP family protein CG45057 |
| **LOC105832294** | -2.01 | 3.58E-22 | 1.82E-20 | Phospholipase A1 |
| **LOC105831727** | -1.29 | 3.82E-22 | 1.93E-20 | Ras-related protein Rab-3 |
| **LOC105836444** | -1.16 | 2.31E-21 | 1.11E-19 | Neurotrimin |
| **LOC105831787** | -1.11 | 3.56E-20 | 1.51E-18 | Zinc finger protein 362 |
| **LOC105835899** | -1.35 | 5.20E-19 | 1.95E-17 | CUGBP Elav-like family member 4 |
| **LOC105835996** | -1.03 | 2.32E-18 | 8.36E-17 | Glutamate-gated chloride channel |
| **LOC105831567** | -1.61 | 8.58E-18 | 2.89E-16 | Transcription factor 21 |
| **LOC105832152** | -1.73 | 1.72E-17 | 5.59E-16 | Esterase E4 |
| **LOC105837623** | -1.26 | 5.68E-17 | 1.71E-15 | Lachesin |
| **LOC105832861** | -1.28 | 9.58E-17 | 2.77E-15 | - |
| **LOC105837307** | -0.89 | 1.00E-16 | 2.89E-15 | Ankyrin-2 |
| **LOC105833207** | -1.39 | 2.45E-16 | 6.83E-15 | Pikachurin |
| **LOC105829008** | -4.29 | 8.32E-16 | 2.21E-14 | - |
| **LOC105830234** | -2.91 | 8.34E-16 | 2.21E-14 | - |
| **LOC105830319** | -4.11 | 8.73E-16 | 2.30E-14 | - |

**Table S3.** Top 20 reproductive-upregulated genes, sorted by FDR. Positive values of Log2 fold change indicate higher expression in reproductive samples.

| Gene | Log2 Fold Change | P-Value | FDR | Description (SwissProt) |
| --- | --- | --- | --- | --- |
| **LOC105837393** | 5.96 | 1.63E-143 | 1.79E-139 | - |
| **LOC105834006** | 4.27 | 7.02E-106 | 3.85E-102 | Gephyrin |
| **LOC105830579** | 3.92 | 1.02E-95 | 3.72E-92 | Cytoplasmic polyadenylation element-binding protein 1 |
| **LOC105834706** | 5.08 | 1.76E-89 | 4.83E-86 | Maternal effect protein oskar |
| **LOC105835926** | 7.37 | 4.12E-87 | 9.03E-84 | Vitellogenin-2 |
| **LOC105840630** | 10.61 | 4.49E-79 | 8.21E-76 | - |
| **LOC105840094** | 4.83 | 3.34E-71 | 5.24E-68 | RCC1 and BTB domain-containing protein 1 |
| **LOC105831415** | 2.52 | 3.38E-68 | 4.64E-65 | - |
| **LOC105834586** | 2.56 | 2.40E-67 | 2.92E-64 | Putative bifunctional UDP-N-acetylglucosamine transferase and deubiquitinase ALG13 |
| **LOC105828810** | 2.77 | 5.75E-65 | 6.30E-62 | Gephyrin |
| **LOC105829254** | 2.81 | 2.21E-63 | 2.20E-60 | Maternal protein exuperantia |
| **LOC105832526** | 2.21 | 1.97E-62 | 1.80E-59 | Protein aubergine |
| **LOC105830728** | 3.29 | 5.56E-62 | 4.69E-59 | S-phase kinase-associated protein 2 |
| **LOC105833392** | 4.08 | 7.20E-62 | 5.64E-59 | Poly(A) RNA polymerase gld-2 homolog A |
| **LOC105828865** | 3.28 | 7.20E-60 | 5.27E-57 | Hyaluronan mediated motility receptor |
| **LOC105837552** | 3.97 | 6.20E-58 | 4.25E-55 | Ribonuclease H1 |
| **LOC105829518** | 2.18 | 1.06E-57 | 6.81E-55 | - |
| **LOC105833500** | 2.19 | 9.12E-57 | 5.56E-54 | Serine/threonine-protein kinase Chk2 |
| **LOC105838098** | 2.29 | 1.10E-56 | 6.37E-54 | - |
| **LOC105833023** | 3.33 | 1.49E-56 | 8.19E-54 | - |

**Table S4.** Summary of the raw phylostrata identified for genes in the *M. pharoanis* genome (Fig. S6), and 6 categories that phylostrata were grouped into “Condensed PS1”. For some analyses that required ∼100 genes in each caste-associated category, we also created a third grouping, “Condensed PS2” that combined the hymenopteran and ant categories.

| **PS** | **Condensed PS1** | **Condensed PS2** |
| --- | --- | --- |
| cellular organisms | cellular | cellular |
| Eukaryota | eukaryote | eukaryote |
| Opisthokonta | bilaterian | bilaterian |
| Metazoa | bilaterian | bilaterian |
| Eumetazoa | bilaterian | bilaterian |
| Bilateria | bilaterian | bilaterian |
| Protostomia | bilaterian | bilaterian |
| Ecdysozoa | insect | insect |
| Arthropoda | insect | insect |
| Pancrustacea | insect | insect |
| Neoptera | insect | insect |
| Endopterygota | insect | insect |
| Apocrita | hymenopteran | hymenopteran |
| Aculeata | hymenopteran | hymenopteran |
| Vespoidea | hymenopteran | hymenopteran |
| Formicidae | ant | hymenopteran |
| Myrmicinae | ant | hymenopteran |
| Solenopsidini | ant | hymenopteran |
| *Monomorium pharaonis* | ant | hymenopteran |

**Table S5.** Top 3 GO terms for workers and reproductives for each differential expression test, as calculated using the R package GOstats, sorted by p-value. L2 not included due to paucity of differentially expressed genes.

| GOBPID | Pvalue | OddsRatio | ExpCount | Count | Size | Term | Sample | Caste |
| --- | --- | --- | --- | --- | --- | --- | --- | --- |
| GO:0006811 | 1.95E-12 | 3.554038863 | 26.99482536 | 63 | 154 | ion transport | MainEffect | Worker |
| GO:0046034 | 3.35E-10 | 20.61047619 | 3.681112549 | 17 | 21 | ATP metabolic process | MainEffect | Worker |
| GO:0007186 | 6.95E-10 | 3.66109831 | 19.10672704 | 46 | 109 | G-protein coupled receptor signaling pathway | MainEffect | Worker |
| GO:0046483 | 4.38E-29 | 2.926335878 | 172.740621 | 287 | 567 | heterocycle metabolic process | MainEffect | Reproductive |
| GO:0090304 | 1.36E-28 | 3.148458605 | 141.0562743 | 246 | 463 | nucleic acid metabolic process | MainEffect | Reproductive |
| GO:1901360 | 2.19E-28 | 2.8789485 | 173.9592497 | 287 | 571 | organic cyclic compound metabolic process | MainEffect | Reproductive |
| GO:0015711 | 0.000725545 | 5.986786787 | 1.629366106 | 7 | 22 | organic anion transport | LarvalMain | Worker |
| GO:0015849 | 0.000725545 | 5.986786787 | 1.629366106 | 7 | 22 | organic acid transport | LarvalMain | Worker |
| GO:0046942 | 0.000725545 | 5.986786787 | 1.629366106 | 7 | 22 | carboxylic acid transport | LarvalMain | Worker |
| GO:1902578 | 1.28E-05 | 2.016145186 | 36.39068564 | 61 | 388 | single-organism localization | LarvalMain | Reproductive |
| GO:0044699 | 1.45E-05 | 1.69630845 | 140.6856404 | 175 | 1500 | single-organism process | LarvalMain | Reproductive |
| GO:0044765 | 1.58E-05 | 2.009181745 | 35.82794308 | 60 | 382 | single-organism transport | LarvalMain | Reproductive |
| GO:0006040 | 1.24E-06 | 7.450946644 | 2.302716688 | 12 | 40 | amino sugar metabolic process | L3 | Worker |
| GO:0006030 | 1.24E-06 | 7.450946644 | 2.302716688 | 12 | 40 | chitin metabolic process | L3 | Worker |
| GO:1901071 | 1.24E-06 | 7.450946644 | 2.302716688 | 12 | 40 | glucosamine-containing compound metabolic process | L3 | Worker |
| GO:0006720 | 1.96E-05 | 14.98484848 | 0.626778784 | 6 | 17 | isoprenoid metabolic process | L3 | Reproductive |
| GO:0008299 | 1.96E-05 | 14.98484848 | 0.626778784 | 6 | 17 | isoprenoid biosynthetic process | L3 | Reproductive |
| GO:0044255 | 0.000359857 | 4.730458221 | 2.285899094 | 9 | 62 | cellular lipid metabolic process | L3 | Reproductive |
| GO:0007600 | 7.99E-12 | 5.110028653 | 11.4851229 | 37 | 92 | sensory perception | L4 | Worker |
| GO:0007606 | 7.99E-12 | 5.110028653 | 11.4851229 | 37 | 92 | sensory perception of chemical stimulus | L4 | Worker |
| GO:0003008 | 7.99E-12 | 5.110028653 | 11.4851229 | 37 | 92 | system process | L4 | Worker |
| GO:0044710 | 1.73E-09 | 2.068259836 | 82.89844761 | 130 | 671 | single-organism metabolic process | L4 | Reproductive |
| GO:0055114 | 1.23E-08 | 2.372507113 | 40.52263907 | 75 | 328 | oxidation-reduction process | L4 | Reproductive |
| **GO:0005975** | 9.96E-06 | 2.690731282 | 15.44307891 | 33 | 125 | carbohydrate metabolic process | L4 | Reproductive |
| **GO:0007608** | 3.27E-09 | 6.209081836 | 6.119340233 | 23 | 53 | sensory perception of smell | L5 | Worker |
| **GO:0007600** | 6.01E-07 | 3.551861702 | 10.62225097 | 28 | 92 | sensory perception | L5 | Worker |
| **GO:0007606** | 6.01E-07 | 3.551861702 | 10.62225097 | 28 | 92 | sensory perception of chemical stimulus | L5 | Worker |
| **GO:0044710** | 2.09E-08 | 1.99925 | 78.3412031 | 121 | 671 | single-organism metabolic process | L5 | Reproductive |
| **GO:0055114** | 7.45E-08 | 2.305735369 | 38.29495472 | 70 | 328 | oxidation-reduction process | L5 | Reproductive |
| **GO:0008152** | 0.000452937 | 1.509656659 | 233.6225744 | 262 | 2001 | metabolic process | L5 | Reproductive |
| **GO:0055114** | 1.70E-26 | 3.543282815 | 119.3402329 | 209 | 328 | oxidation-reduction process | Gaster | Worker |
| **GO:0044699** | 7.12E-15 | 1.788020026 | 545.76326 | 649 | 1500 | single-organism process | Gaster | Worker |
| **GO:0055085** | 3.98E-14 | 2.877425945 | 81.50064683 | 135 | 224 | transmembrane transport | Gaster | Worker |
| **GO:0044260** | 7.56E-57 | 3.762404675 | 279.5653299 | 469 | 952 | cellular macromolecule metabolic process | Gaster | Reproductive |
| **GO:0090304** | 1.66E-56 | 5.213744618 | 135.9650712 | 286 | 463 | nucleic acid metabolic process | Gaster | Reproductive |
| **GO:0006139** | 1.18E-48 | 4.196444744 | 159.7516171 | 307 | 544 | nucleobase-containing compound metabolic process | Gaster | Reproductive |
| **GO:0046034** | 2.31E-10 | 21.1920078 | 3.599611902 | 17 | 21 | ATP metabolic process | Head | Worker |
| **GO:0006163** | 2.16E-09 | 7.100674262 | 7.027813713 | 24 | 41 | purine nucleotide metabolic process | Head | Worker |
| **GO:0009161** | 2.77E-09 | 14.11695906 | 3.942432083 | 17 | 23 | ribonucleoside monophosphate metabolic process | Head | Worker |
| **GO:0006259** | 9.13E-09 | 3.252254768 | 22.91332471 | 49 | 108 | DNA metabolic process | Head | Reproductive |
| **GO:0044710** | 4.50E-07 | 1.657178803 | 142.3596378 | 190 | 671 | single-organism metabolic process | Head | Reproductive |
| **GO:0006950** | 1.40E-06 | 3.418803419 | 14.42690815 | 32 | 68 | response to stress | Head | Reproductive |

**Table S6.** Top 3 GO terms for each phylostrata category for each differential expression test, as calculated using the R package GOstats, sorted by p-value. L2 not included due to paucity of differentially expressed genes. Missing phylostrata categories returned no significant GO terms.

| GOBPID | Pvalue | OddsRatio | ExpCount | Count | Size | Term | Sample | Caste | Phylostrata |
| --- | --- | --- | --- | --- | --- | --- | --- | --- | --- |
| GO:0006811 | 6.23E-12 | 4.075406342 | 16.13712807 | 46 | 154 | ion transport | MainEffect | Worker | cellular |
| GO:0044765 | 2.17E-11 | 2.73200443 | 40.02846054 | 81 | 382 | single-organism transport | MainEffect | Worker | cellular |
| GO:0055085 | 2.20E-11 | 3.324968041 | 23.47218629 | 57 | 224 | transmembrane transport | MainEffect | Worker | cellular |
| GO:0065007 | 1.16E-06 | 3.287946429 | 14.76746442 | 33 | 593 | biological regulation | MainEffect | Worker | eukaryote |
| GO:0050789 | 2.40E-06 | 3.19417122 | 14.46862872 | 32 | 581 | regulation of biological process | MainEffect | Worker | eukaryote |
| GO:0050794 | 5.66E-06 | 3.074883684 | 14.26940492 | 31 | 573 | regulation of cellular process | MainEffect | Worker | eukaryote |
| GO:0007186 | 1.46E-26 | 21.5191793 | 3.243208279 | 33 | 109 | G-protein coupled receptor signaling pathway | MainEffect | Worker | bilaterian |
| GO:0050794 | 1.41E-23 | 9.089912281 | 17.04915912 | 60 | 573 | regulation of cellular process | MainEffect | Worker | bilaterian |
| GO:0050789 | 3.06E-23 | 8.921545106 | 17.28719276 | 60 | 581 | regulation of biological process | MainEffect | Worker | bilaterian |
| GO:0006040 | 6.78E-05 | 61.78378378 | 0.090556274 | 3 | 40 | amino sugar metabolic process | MainEffect | Worker | insect |
| GO:0006030 | 6.78E-05 | 61.78378378 | 0.090556274 | 3 | 40 | chitin metabolic process | MainEffect | Worker | insect |
| GO:1901071 | 6.78E-05 | 61.78378378 | 0.090556274 | 3 | 40 | glucosamine-containing compound metabolic process | MainEffect | Worker | insect |
| GO:0007600 | 5.91E-39 | 152.978836 | 1.130659767 | 29 | 92 | sensory perception | MainEffect | Worker | hymenopteran_ant |
| GO:0007606 | 5.91E-39 | 152.978836 | 1.130659767 | 29 | 92 | sensory perception of chemical stimulus | MainEffect | Worker | hymenopteran_ant |
| GO:0003008 | 5.91E-39 | 152.978836 | 1.130659767 | 29 | 92 | system process | MainEffect | Worker | hymenopteran_ant |
| GO:0008152 | 7.43E-29 | 3.538233934 | 349.4631307 | 456 | 2001 | metabolic process | MainEffect | Reproductive | cellular |
| GO:0044238 | 3.11E-15 | 2.120088457 | 250.4398448 | 333 | 1434 | primary metabolic process | MainEffect | Reproductive | cellular |
| GO:0071704 | 1.53E-13 | 2.023948498 | 262.839586 | 340 | 1505 | organic substance metabolic process | MainEffect | Reproductive | cellular |
| **GO:0090304** | 9.66E-15 | 3.036420958 | 45.0721216 | 95 | 463 | nucleic acid metabolic process | MainEffect | Reproductive | eukaryote |
| **GO:0051649** | 2.92E-12 | 5.605050139 | 9.345407503 | 34 | 96 | establishment of localization in cell | MainEffect | Reproductive | eukaryote |
| **GO:0051641** | 7.52E-12 | 5.190694127 | 10.12419146 | 35 | 104 | cellular localization | MainEffect | Reproductive | eukaryote |
| **GO:0031323** | 3.77E-17 | 9.529559748 | 5.68305304 | 32 | 191 | regulation of cellular metabolic process | MainEffect | Reproductive | bilaterian |
| **GO:0019222** | 1.56E-16 | 8.99047619 | 5.95084088 | 32 | 200 | regulation of metabolic process | MainEffect | Reproductive | bilaterian |
| **GO:0060255** | 2.58E-16 | 9.141108114 | 5.623544631 | 31 | 189 | regulation of macromolecule metabolic process | MainEffect | Reproductive | bilaterian |
| **GO:0007600** | 5.64E-10 | 123.4470588 | 0.26778784 | 7 | 92 | sensory perception | MainEffect | Reproductive | hymenopteran_ant |
| **GO:0007606** | 5.64E-10 | 123.4470588 | 0.26778784 | 7 | 92 | sensory perception of chemical stimulus | MainEffect | Reproductive | hymenopteran_ant |
| **GO:0003008** | 5.64E-10 | 123.4470588 | 0.26778784 | 7 | 92 | system process | MainEffect | Reproductive | hymenopteran_ant |
| **GO:0006811** | 1.39E-05 | 3.35645933 | 7.620310479 | 21 | 154 | ion transport | LarvalMain | Worker | cellular |
| **GO:0006508** | 0.000148191 | 2.454297821 | 13.01390686 | 27 | 263 | proteolysis | LarvalMain | Worker | cellular |
| **GO:0006820** | 0.000277076 | 5.735960591 | 1.781371281 | 8 | 36 | anion transport | LarvalMain | Worker | cellular |
| **GO:0065007** | 0.000350266 | 4.004145078 | 5.561772316 | 14 | 593 | biological regulation | LarvalMain | Worker | eukaryote |
| **GO:0007165** | 0.000783482 | 4.257379353 | 3.254527814 | 10 | 347 | signal transduction | LarvalMain | Worker | eukaryote |
| **GO:0044700** | 0.000820369 | 4.229156963 | 3.273285899 | 10 | 349 | single organism signaling | LarvalMain | Worker | eukaryote |
| **GO:0050794** | 4.20E-12 | 10.33104396 | 7.227360931 | 27 | 573 | regulation of cellular process | LarvalMain | Worker | bilaterian |
| **GO:0050789** | 5.93E-12 | 10.14936823 | 7.328266494 | 27 | 581 | regulation of biological process | LarvalMain | Worker | bilaterian |
| **GO:0065007** | 9.88E-12 | 9.886484099 | 7.479624838 | 27 | 593 | biological regulation | LarvalMain | Worker | bilaterian |
| **GO:0006040** | 0.025709<br>996 | 78.23076923 | 0.025873221 | 1 | 40 | amino sugar metabolic process | LarvalMain | Worker | insect |
| **GO:0006030** | 0.025709<br>996 | 78.23076923 | 0.025873221 | 1 | 40 | chitin metabolic process | LarvalMain | Worker | insect |
| **GO:1901071** | 0.025709<br>996 | 78.23076923 | 0.025873221 | 1 | 40 | glucosamine-containing<br>compound metabolic process | LarvalMain | Worker | insect |
| **GO:0055085** | 1.46E-06 | 2.87593985 | 14.63389392 | 34 | 224 | transmembrane transport | LarvalMain | Reproductive | cellular |
| **GO:0044710** | 3.85E-06 | 2.068575064 | 43.83635188 | 71 | 671 | single-organism metabolic<br>process | LarvalMain | Reproductive | cellular |
| **GO:0055114** | 9.66E-05 | 2.153374233 | 21.42820181 | 39 | 328 | oxidation-reduction process | LarvalMain | Reproductive | cellular |
| **GO:0006281** | 2.68E-05 | 9.741395349 | 0.921733506 | 7 | 57 | DNA repair | LarvalMain | Reproductive | eukaryote |
| **GO:0006974** | 3.38E-05 | 9.360465116 | 0.954075032 | 7 | 59 | cellular response to DNA damage<br>stimulus | LarvalMain | Reproductive | eukaryote |
| **GO:0033554** | 3.78E-05 | 9.180781044 | 0.970245796 | 7 | 60 | cellular response to stress | LarvalMain | Reproductive | eukaryote |
| **GO:0006869** | 3.01E-11 | 111.3072917 | 0.150388098 | 7 | 15 | lipid transport | LarvalMain | Reproductive | bilaterian |
| **GO:0010876** | 3.01E-11 | 111.3072917 | 0.150388098 | 7 | 15 | lipid localization | LarvalMain | Reproductive | bilaterian |
| **GO:0033036** | 1.54E-05 | 11.00949367 | 0.862225097 | 7 | 86 | macromolecule localization | LarvalMain | Reproductive | bilaterian |
| **GO:0007600** | 2.40E-05 | 45.40909091 | 0.208279431 | 4 | 92 | sensory perception | LarvalMain | Reproductive | hymenopteran<br>_ant |
| **GO:0007606** | 2.40E-05 | 45.40909091 | 0.208279431 | 4 | 92 | sensory perception of chemical<br>stimulus | LarvalMain | Reproductive | hymenopteran<br>_ant |
| **GO:0003008** | 2.40E-05 | 45.40909091 | 0.208279431 | 4 | 92 | system process | LarvalMain | Reproductive | hymenopteran<br>_ant |
| **GO:0044765** | 6.83E-06 | 2.752626552 | 15.19598965 | 33 | 382 | single-organism transport | L3 | Worker | cellular |
| **GO:1902578** | 9.64E-06 | 2.699906103 | 15.43467012 | 33 | 388 | single-organism localization | L3 | Worker | cellular |
| **GO:0055114** | 1.86E-05 | 2.754927773 | 13.04786546 | 29 | 328 | oxidation-reduction process | L3 | Worker | cellular |
| **GO:0030328** | 0.004204398 | Inf | 0.004204398 | 1 | 1 | prenylcysteine catabolic process | L3 | Worker | eukaryote |
| **GO:0000098** | 0.004204398 | Inf | 0.004204398 | 1 | 1 | sulfur amino acid catabolic process | L3 | Worker | eukaryote |
| **GO:0030329** | 0.004204398 | Inf | 0.004204398 | 1 | 1 | prenylcysteine metabolic process | L3 | Worker | eukaryote |
| **GO:0007186** | 4.68E-08 | 14.24723425 | 1.092820181 | 10 | 109 | G-protein coupled receptor signaling pathway | L3 | Worker | bilaterian |
| **GO:0050794** | 9.65E-07 | 6.251975052 | 5.744825356 | 18 | 573 | regulation of cellular process | L3 | Worker | bilaterian |
| **GO:0050789** | 1.20E-06 | 6.143462222 | 5.825032342 | 18 | 581 | regulation of biological process | L3 | Worker | bilaterian |
| **GO:0006040** | 5.78E-09 | 217.8571429 | 0.090556274 | 5 | 40 | amino sugar metabolic process | L3 | Worker | insect |
| **GO:0006030** | 5.78E-09 | 217.8571429 | 0.090556274 | 5 | 40 | chitin metabolic process | L3 | Worker | insect |
| **GO:1901071** | 5.78E-09 | 217.8571429 | 0.090556274 | 5 | 40 | glucosamine-containing compound metabolic process | L3 | Worker | insect |
| **GO:0050909** | 0.012613195 | Inf | 0.012613195 | 1 | 39 | sensory perception of taste | L3 | Worker | hymenopteran_ant |
| **GO:0007600** | 0.029754204 | Inf | 0.029754204 | 1 | 92 | sensory perception | L3 | Worker | hymenopteran_ant |
| **GO:0007606** | 0.029754204 | Inf | 0.029754204 | 1 | 92 | sensory perception of chemical stimulus | L3 | Worker | hymenopteran_ant |
| **GO:0006720** | 4.38E-05 | 17.1347032 | 0.428848642 | 5 | 17 | isoprenoid metabolic process | L3 | Reproductive | cellular |
| **GO:0008299** | 4.38E-05 | 17.1347032 | 0.428848642 | 5 | 17 | isoprenoid biosynthetic process | L3 | Reproductive | cellular |
| **GO:0044255** | 0.000128384 | 6.264550265 | 1.564036223 | 8 | 62 | cellular lipid metabolic process | L3 | Reproductive | cellular |
| **GO:0010970** | 0.005174644 | Inf | 0.005174644 | 1 | 1 | establishment of localization by movement along microtubule | L3 | Reproductive | eukaryote |
| **GO:0098840** | 0.005174644 | Inf | 0.005174644 | 1 | 1 | protein transport along microtubule | L3 | Reproductive | eukaryote |
| **GO:0042073** | 0.005174644 | Inf | 0.005174644 | 1 | 1 | intraciliary transport | L3 | Reproductive | eukaryote |
| **GO:0050794** | 9.40E-06 | 20.08244681 | 2.038486417 | 9 | 573 | regulation of cellular process | L3 | Reproductive | bilaterian |
| **GO:0050789** | 1.06E-05 | 19.73863636 | 2.06694696 | 9 | 581 | regulation of biological process | L3 | Reproductive | bilaterian |
| **GO:0065007** | 1.26E-05 | 19.24058219 | 2.109637775 | 9 | 593 | biological regulation | L3 | Reproductive | bilaterian |
| **GO:0007600** | 2.40E-06 | 43.04597701 | 0.26778784 | 5 | 92 | sensory perception | L3 | Reproductive | hymenopteran_ant |
| **GO:0007606** | 2.40E-06 | 43.04597701 | 0.26778784 | 5 | 92 | sensory perception of chemical stimulus | L3 | Reproductive | hymenopteran_ant |
| **GO:0003008** | 2.40E-06 | 43.04597701 | 0.26778784 | 5 | 92 | system process | L3 | Reproductive | hymenopteran_ant |
| **GO:0006508** | 7.18E-09 | 3.162303099 | 18.11739974 | 44 | 263 | proteolysis | L4 | Worker | cellular |
| **GO:0006811** | 3.41E-06 | 3.145542291 | 10.60866753 | 27 | 154 | ion transport | L4 | Worker | cellular |
| **GO:0055114** | 3.00E-05 | 2.226843332 | 22.59508409 | 42 | 328 | oxidation-reduction process | L4 | Worker | cellular |
| **GO:0065007** | 9.05E-11 | 5.535394265 | 12.08247089 | 35 | 593 | biological regulation | L4 | Worker | eukaryote |
| **GO:0050789** | 1.43E-09 | 4.980109489 | 11.83796895 | 33 | 581 | regulation of biological process | L4 | Worker | eukaryote |
| **GO:0050794** | 4.85E-09 | 4.747242263 | 11.67496766 | 32 | 573 | regulation of cellular process | L4 | Worker | eukaryote |
| **GO:0050794** | 4.03E-19 | 11.27285115 | 11.30433376 | 43 | 573 | regulation of cellular process | L4 | Worker | bilaterian |
| **GO:0050789** | 7.08E-19 | 11.0697026 | 11.46216041 | 43 | 581 | regulation of biological process | L4 | Worker | bilaterian |
| **GO:0065007** | 1.62E-18 | 10.77606061 | 11.69890039 | 43 | 593 | biological regulation | L4 | Worker | bilaterian |
| **GO:0007600** | 1.37E-51 | 274.8673469 | 1.279430789 | 36 | 92 | sensory perception | L4 | Worker | hymenopteran_ant |
| **GO:0007606** | 1.37E-51 | 274.8673469 | 1.279430789 | 36 | 92 | sensory perception of chemical stimulus | L4 | Worker | hymenopteran_ant |
| **GO:0003008** | 1.37E-51 | 274.8673469 | 1.279430789 | 36 | 92 | system process | L4 | Worker | hymenopteran_ant |
| **GO:0044710** | 2.90E-13 | 2.638315484 | 63.1503881 | 115 | 671 | single-organism metabolic process | L4 | Reproductive | cellular |
| **GO:0055114** | 2.52E-12 | 3.121996562 | 30.86934023 | 70 | 328 | oxidation-reduction process | L4 | Reproductive | cellular |
| **GO:0008152** | 3.63E-09 | 2.293970547 | 188.3217982 | 232 | 2001 | metabolic process | L4 | Reproductive | cellular |
| **GO:0006810** | 0.002909<br>657 | 2.793736501 | 6.479301423 | 14 | 477 | transport | L4 | Reproductive | eukaryote |
| **GO:0051234** | 0.003089<br>857 | 2.772532189 | 6.520051746 | 14 | 480 | establishment of localization | L4 | Reproductive | eukaryote |
| **GO:0051179** | 0.003687<br>396 | 2.710526316 | 6.642302717 | 14 | 489 | localization | L4 | Reproductive | eukaryote |
| **GO:0006869** | 1.81E-12 | 112.2949309 | 0.18919793 | 8 | 15 | lipid transport | L4 | Reproductive | bilaterian |
| **GO:0010876** | 1.81E-12 | 112.2949309 | 0.18919793 | 8 | 15 | lipid localization | L4 | Reproductive | bilaterian |
| **GO:0033036** | 7.86E-06 | 9.842845327 | 1.084734799 | 8 | 86 | macromolecule localization | L4 | Reproductive | bilaterian |
| **GO:0006040** | 1.19E-07 | 339 | 0.064683053 | 4 | 40 | amino sugar metabolic process | L4 | Reproductive | insect |
| **GO:0006030** | 1.19E-07 | 339 | 0.064683053 | 4 | 40 | chitin metabolic process | L4 | Reproductive | insect |
| **GO:1901071** | 1.19E-07 | 339 | 0.064683053 | 4 | 40 | glucosamine-containing<br>compound metabolic process | L4 | Reproductive | insect |
| **GO:0007600** | 9.98E-05 | 101.0898876 | 0.119016818 | 3 | 92 | sensory perception | L4 | Reproductive | hymenopteran<br>_ant |
| **GO:0007606** | 9.98E-05 | 101.0898876 | 0.119016818 | 3 | 92 | sensory perception of chemical<br>stimulus | L4 | Reproductive | hymenopteran<br>_ant |
| **GO:0003008** | 9.98E-05 | 101.0898876 | 0.119016818 | 3 | 92 | system process | L4 | Reproductive | hymenopteran<br>_ant |
| **GO:0055114** | 4.19E-08 | 2.723857659 | 23.97412678 | 51 | 328 | oxidation-reduction process | L5 | Worker | cellular |
| **GO:0006508** | 7.65E-06 | 2.459845302 | 19.22315653 | 39 | 263 | proteolysis | L5 | Worker | cellular |
| **GO:0006811** | 8.94E-05 | 2.638937097 | 11.25614489 | 25 | 154 | ion transport | L5 | Worker | cellular |
| **GO:0065007** | 2.52E-10 | 6.7114595 | 9.205692109 | 29 | 593 | biological regulation | L5 | Worker | eukaryote |
| **GO:0050789** | 9.91E-10 | 6.306329114 | 9.019404916 | 28 | 581 | regulation of biological process | L5 | Worker | eukaryote |
| **GO:0050794** | 4.39E-09 | 5.882260597 | 8.895213454 | 27 | 573 | regulation of cellular process | L5 | Worker | eukaryote |
| **GO:0050794** | 1.83E-13 | 10.65023374 | 7.968628719 | 30 | 573 | regulation of cellular process | L5 | Worker | bilaterian |
| **GO:0050789** | 2.70E-13 | 10.46209689 | 8.079883571 | 30 | 581 | regulation of biological process | L5 | Worker | bilaterian |
| **GO:0065007** | 4.77E-13 | 10.18991666 | 8.246765847 | 30 | 593 | biological regulation | L5 | Worker | bilaterian |
| **GO:0006040** | 0.025709<br>996 | 78.23076923 | 0.025873221 | 1 | 40 | amino sugar metabolic process | L5 | Worker | insect |
| **GO:0006030** | 0.025709<br>996 | 78.23076923 | 0.025873221 | 1 | 40 | chitin metabolic process | L5 | Worker | insect |
| **GO:1901071** | 0.025709<br>996 | 78.23076923 | 0.025873221 | 1 | 40 | glucosamine-containing<br>compound metabolic process | L5 | Worker | insect |
| **GO:0007600** | 4.56E-37 | 177.6065934 | 1.01164295 | 27 | 92 | sensory perception | L5 | Worker | hymenopteran<br>_ant |
| **GO:0007606** | 4.56E-37 | 177.6065934 | 1.01164295 | 27 | 92 | sensory perception of chemical<br>stimulus | L5 | Worker | hymenopteran<br>_ant |
| **GO:0003008** | 4.56E-37 | 177.6065934 | 1.01164295 | 27 | 92 | system process | L5 | Worker | hymenopteran<br>_ant |
| **GO:0044710** | 2.40E-13 | 2.754534915 | 57.0740621 | 107 | 671 | single-organism metabolic<br>process | L5 | Reprodu<br>ctive | cellular |
| **GO:0055114** | 4.46E-13 | 3.363359137 | 27.89909444 | 67 | 328 | oxidation-reduction process | L5 | Reprodu<br>ctive | cellular |
| **GO:0008152** | 1.75E-08 | 2.296387598 | 170.2014877 | 210 | 200<br>1 | metabolic process | L5 | Reprodu<br>ctive | cellular |
| **GO:0051649** | 0.000343<br>296 | 5.356363636 | 1.800776197 | 8 | 96 | establishment of localization in<br>cell | L5 | Reprodu<br>ctive | eukaryote |
| **GO:0022406** | 0.000485<br>846 | 27.52727273 | 0.168822768 | 3 | 9 | membrane docking | L5 | Reprodu<br>ctive | eukaryote |
| **GO:0051641** | 0.000593<br>227 | 4.896666667 | 1.95084088 | 8 | 104 | cellular localization | L5 | Reprodu<br>ctive | eukaryote |
| **GO:0006869** | 1.05E-05 | 41.07744108 | 0.150388098 | 4 | 15 | lipid transport | L5 | Reprodu<br>ctive | bilaterian |
| **GO:0010876** | 1.05E-05 | 41.07744108 | 0.150388098 | 4 | 15 | lipid localization | L5 | Reprodu<br>ctive | bilaterian |
| **GO:0050794** | 0.000569<br>083 | 3.685993897 | 5.744825356 | 14 | 573 | regulation of cellular process | L5 | Reprodu<br>ctive | bilaterian |
| **GO:0006040** | 2.01E-06 | Inf | 0.038809832 | 3 | 40 | amino sugar metabolic process | L5 | Reprodu<br>ctive | insect |
| **GO:0006030** | 2.01E-06 | Inf | 0.038809832 | 3 | 40 | chitin metabolic process | L5 | Reproductive | insect |
| **GO:1901071** | 2.01E-06 | Inf | 0.038809832 | 3 | 40 | glucosamine-containing compound metabolic process | L5 | Reproductive | insect |
| **GO:0007600** | 0.000244237 | 50.52808989 | 0.148771022 | 3 | 92 | sensory perception | L5 | Reproductive | hymenopteran_ant |
| **GO:0007606** | 0.000244237 | 50.52808989 | 0.148771022 | 3 | 92 | sensory perception of chemical stimulus | L5 | Reproductive | hymenopteran_ant |
| **GO:0003008** | 0.000244237 | 50.52808989 | 0.148771022 | 3 | 92 | system process | L5 | Reproductive | hymenopteran_ant |
| **GO:0055114** | 5.29E-45 | 5.51156392 | 81.57567917 | 194 | 328 | oxidation-reduction process | Gaster | Worker | cellular |
| **GO:0044710** | 3.59E-28 | 2.829303501 | 166.8819534 | 280 | 671 | single-organism metabolic process | Gaster | Worker | cellular |
| **GO:0055085** | 7.96E-18 | 3.432707097 | 55.71021992 | 113 | 224 | transmembrane transport | Gaster | Worker | cellular |
| **GO:0015991** | 5.34E-07 | 44.71212121 | 0.401681759 | 6 | 9 | ATP hydrolysis coupled proton transport | Gaster | Worker | eukaryote |
| **GO:0015988** | 5.34E-07 | 44.71212121 | 0.401681759 | 6 | 9 | energy coupled proton transmembrane transport, against electrochemical gradient | Gaster | Worker | eukaryote |
| **GO:0090662** | 5.34E-07 | 44.71212121 | 0.401681759 | 6 | 9 | ATP hydrolysis coupled transmembrane transport | Gaster | Worker | eukaryote |
| **GO:0007186** | 4.14E-34 | 20.91978093 | 5.005821475 | 45 | 109 | G-protein coupled receptor signaling pathway | Gaster | Worker | bilaterian |
| **GO:0050794** | 3.55E-30 | 7.766867116 | 26.31500647 | 86 | 573 | regulation of cellular process | Gaster | Worker | bilaterian |
| **GO:0065007** | 7.35E-30 | 7.640244341 | 27.23350582 | 87 | 593 | biological regulation | Gaster | Worker | bilaterian |
| **GO:0006040** | 3.91E-05 | 82.40540541 | 0.077619664 | 3 | 40 | amino sugar metabolic process | Gaster | Worker | insect |
| **GO:0006030** | 3.91E-05 | 82.40540541 | 0.077619664 | 3 | 40 | chitin metabolic process | Gaster | Worker | insect |
| **GO:1901071** | 3.91E-05 | 82.40540541 | 0.077619664 | 3 | 40 | glucosamine-containing compound metabolic process | Gaster | Worker | insect |
| **GO:0007600** | 2.83E-72 | 265.3066202 | 1.934023286 | 51 | 92 | sensory perception | Gaster | Worker | hymenopteran_ant |
| **GO:0007606** | 2.83E-72 | 265.3066202 | 1.934023286 | 51 | 92 | sensory perception of chemical stimulus | Gaster | Worker | hymenopteran_ant |
| **GO:0003008** | 2.83E-72 | 265.3066202 | 1.934023286 | 51 | 92 | system process | Gaster | Worker | hymenopteran_ant |
| **GO:0044237** | 3.95E-32 | 3.340921765 | 174.6005821 | 290 | 1161 | cellular metabolic process | Gaster | Reproductive | cellular |
| **GO:0044260** | 6.29E-32 | 3.362369338 | 143.1694696 | 255 | 952 | cellular macromolecule metabolic process | Gaster | Reproductive | cellular |
| **GO:0044238** | 2.96E-26 | 2.997337956 | 215.656533 | 320 | 1434 | primary metabolic process | Gaster | Reproductive | cellular |
| **GO:0090304** | 2.39E-24 | 3.851938105 | 52.70892626 | 124 | 463 | nucleic acid metabolic process | Gaster | Reproductive | eukaryote |
| **GO:0016070** | 2.38E-19 | 3.639424649 | 41.55239327 | 99 | 365 | RNA metabolic process | Gaster | Reproductive | eukaryote |
| **GO:0006139** | 4.54E-18 | 3.050319094 | 61.9301423 | 125 | 544 | nucleobase-containing compound metabolic process | Gaster | Reproductive | eukaryote |
| **GO:0050794** | 4.03E-15 | 6.162917195 | 15.3813066 | 47 | 573 | regulation of cellular process | Gaster | Reproductive | bilaterian |
| **GO:0050789** | 7.11E-15 | 6.051029963 | 15.59605433 | 47 | 581 | regulation of biological process | Gaster | Reproductive | bilaterian |
| **GO:0065007** | 1.63E-14 | 5.889346764 | 15.91817594 | 47 | 593 | biological regulation | Gaster | Reproductive | bilaterian |
| **GO:0007591** | 0.000323415 | Inf | 0.000323415 | 1 | 1 | molting cycle, chitin-based cuticle | Gaster | Reproductive | insect |
| **GO:0018990** | 0.000323415 | Inf | 0.000323415 | 1 | 1 | ecdysis, chitin-based cuticle | Gaster | Reproductive | insect |
| **GO:0022404** | 0.000323415 | Inf | 0.000323415 | 1 | 1 | molting cycle process | Gaster | Reproductive | insect |
| **GO:0007608** | 9.18E-05 | 60.72 | 0.102846054 | 3 | 53 | sensory perception of smell | Gaster | Reproductive | hymenopteran_ant |
| **GO:0007600** | 0.000477944 | 33.6741573 | 0.178525226 | 3 | 92 | sensory perception | Gaster | Reproductive | hymenopteran_ant |
| **GO:0007606** | 0.000477944 | 33.6741573 | 0.178525226 | 3 | 92 | sensory perception of chemical stimulus | Gaster | Reproductive | hymenopteran_ant |
| **GO:0044765** | 1.70E-11 | 2.784824172 | 38.42238034 | 79 | 382 | single-organism transport | Head | Worker | cellular |
| **GO:1902578** | 3.97E-11 | 2.724137931 | 39.02587322 | 79 | 388 | single-organism localization | Head | Worker | cellular |
| **GO:0006836** | 3.03E-10 | 37.1638796 | 1.508732212 | 12 | 15 | neurotransmitter transport | Head | Worker | cellular |
| **GO:0050789** | 7.05E-08 | 3.619816514 | 15.2202458 | 36 | 581 | regulation of biological process | Head | Worker | eukaryote |
| **GO:0065007** | 1.22E-07 | 3.52459605 | 15.53460543 | 36 | 593 | biological regulation | Head | Worker | eukaryote |
| **GO:0050794** | 6.18E-07 | 3.317727865 | 15.0106727 | 34 | 573 | regulation of cellular process | Head | Worker | eukaryote |
| **GO:0007186** | 1.59E-27 | 25.41125541 | 2.820181113 | 32 | 109 | G-protein coupled receptor signaling pathway | Head | Worker | bilaterian |
| **GO:0050794** | 2.26E-22 | 9.976433971 | 14.82535576 | 54 | 573 | regulation of cellular process | Head | Worker | bilaterian |
| **GO:0050789** | 4.58E-22 | 9.793460809 | 15.03234153 | 54 | 581 | regulation of biological process | Head | Worker | bilaterian |
| **GO:0006040** | 2.59E-10 | 179.3529412 | 0.116429495 | 6 | 40 | amino sugar metabolic process | Head | Worker | insect |
| **GO:0006030** | 2.59E-10 | 179.3529412 | 0.116429495 | 6 | 40 | chitin metabolic process | Head | Worker | insect |
| **GO:1901071** | 2.59E-10 | 179.3529412 | 0.116429495 | 6 | 40 | glucosamine-containing compound metabolic process | Head | Worker | insect |
| **GO:0007600** | 3.24E-49 | 229.6491228 | 1.279430789 | 35 | 92 | sensory perception | Head | Worker | hymenopteran_ant |
| **GO:0007606** | 3.24E-49 | 229.6491228 | 1.279430789 | 35 | 92 | sensory perception of chemical stimulus | Head | Worker | hymenopteran_ant |
| **GO:0003008** | 3.24E-49 | 229.6491228 | 1.279430789 | 35 | 92 | system process | Head | Worker | hymenopteran_ant |
| **GO:0008152** | 6.79E-27 | 4.180646933 | 262.097348 | 352 | 2001 | metabolic process | Head | Reproductive | cellular |
| **GO:0044710** | 3.33E-15 | 2.509660566 | 87.88971539 | 152 | 671 | single-organism metabolic process | Head | Reproductive | cellular |
| **GO:0055114** | 1.44E-08 | 2.325211009 | 42.96248383 | 78 | 328 | oxidation-reduction process | Head | Reproductive | cellular |
| **GO:0007017** | 1.05E-07 | 8.821647059 | 2.249029754 | 13 | 38 | microtubule-based process | Head | Reproductive | eukaryote |
| **GO:0006259** | 7.08E-07 | 4.204767986 | 6.391979301 | 21 | 108 | DNA metabolic process | Head | Reproductive | eukaryote |
| **GO:0033554** | 1.09E-06 | 5.682539683 | 3.551099612 | 15 | 60 | cellular response to stress | Head | Reproductive | eukaryote |
| **GO:0065007** | 5.29E-08 | 4.252502224 | 12.08247089 | 31 | 593 | biological regulation | Head | Reproductive | bilaterian |
| **GO:0050789** | 1.38E-07 | 4.088434252 | 11.83796895 | 30 | 581 | regulation of biological process | Head | Reproductive | bilaterian |
| **GO:0050794** | 4.08E-07 | 3.896247837 | 11.67496766 | 29 | 573 | regulation of cellular process | Head | Reproductive | bilaterian |
| **GO:0048598** | 0.001293661 | Inf | 0.001293661 | 1 | 1 | embryonic morphogenesis | Head | Reproductive | hymenopteran_ant |
| **GO:0035434** | 0.003877217 | 514.3333333 | 0.003880983 | 1 | 3 | copper ion transmembrane transport | Head | Reproductive | hymenopteran_ant |
| **GO:0009790** | 0.003877217 | 514.3333333 | 0.003880983 | 1 | 3 | embryo development | Head | Reproductive | hymenopteran_ant |

**Table S7.** Model selection parameters from MKtest2.0 (*13*, *14*) for estimating *α*, the proportion of amino acid substitution driven by positive selection. The first three columns show the number of parameters for *α* and *f*, as well as the total number of model parameters, *K*. We mainly considered models with per-class estimates (i.e. three separate estimates for worker-associated, reproductive-associated, and NDE genes) for both *α* and *f*, or with per-locus estimates for *f*. Of these two main models in bold that we considered, the model including per-locus estimates of *f* fit the data much better. We focus on results from this model, although the per-class *α* and *f* model produced very similar results, showing the same pattern and overlapping *α* estimates. We also show results from models where *α* and/or *f* is fixed or had a single, genome-wide estimate. LnL maximized log likelihood; AIC, Akaike information criterion; AICc, second-order AIC; BIC, Bayesian information criterion (Welch 2006; Obbard et al. 2009).

| Model description | $\alpha$ | $f$ | $K$ | $LnL$ | AIC | AICc | BIC |
| --- | --- | --- | --- | --- | --- | --- | --- |
|  | 0 | 0 | 2 | -1002096.22231 | 2004196.44462 | 2004196.44495 | 2004213.43161 |
|  | 3 | 0 | 5 | -678989.00409 | 1357988.00818 | 1357988.00984 | 1358030.4765 |
|  | 1 | 0 | 3 | -615656.376167 | 1231318.75233 | 1231318.753 | 1231344.23282 |
|  | 0 | 1 | 3 | -522404.908371 | 1044815.81674 | 1044815.81741 | 1044841.29722 |
|  | 1 | 1 | 4 | -521956.917522 | 1043921.83504 | 1043921.83615 | 1043955.80902 |
|  | 1 | 3 | 6 | -521585.629106 | 1043183.25821 | 1043183.26054 | 1043234.21918 |
|  | 3 | 1 | 6 | -521394.185871 | 1042800.37174 | 1042800.37407 | 1042851.33271 |
| **Per-class $\alpha$ and $f$** | **3** | **3** | **8** | **-521349.0028** | **1042714.0056** | **1042714.00959** | **1042781.95355** |
|  | 0 | 9020 | 9022 | -378721.983142 | 775487.966284 | 781505.300505 | 852116.268919 |
|  | 1 | 9020 | 9023 | -377650.16441 | 773346.32882 | 779365.219417 | 849983.124949 |
| **Per-class $\alpha$ and per-locus $f$** | **3** | **9020** | **9025** | **-377432.859261** | **772917.718522** | **778937/722662** | **849569.501639** |

**Table S8.**
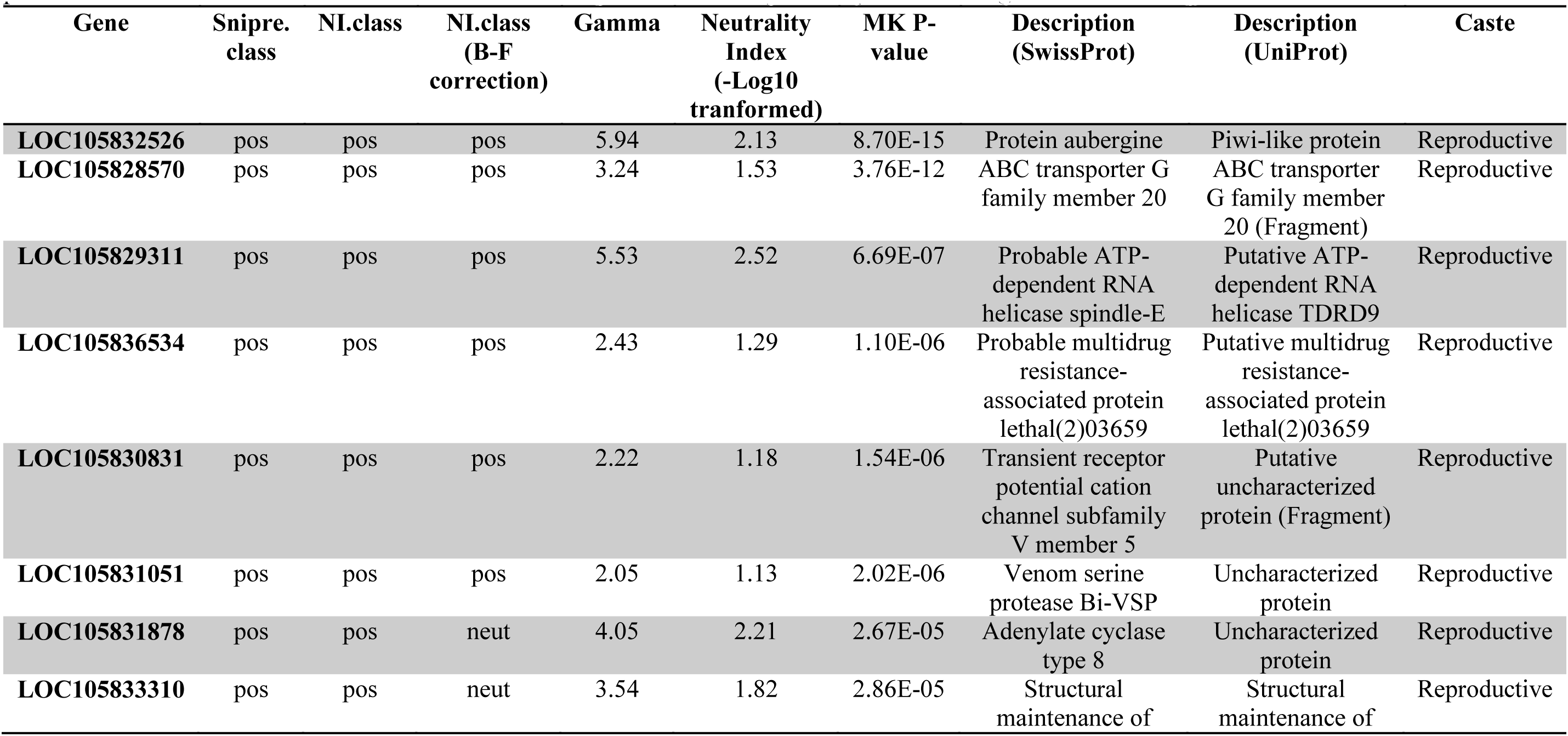
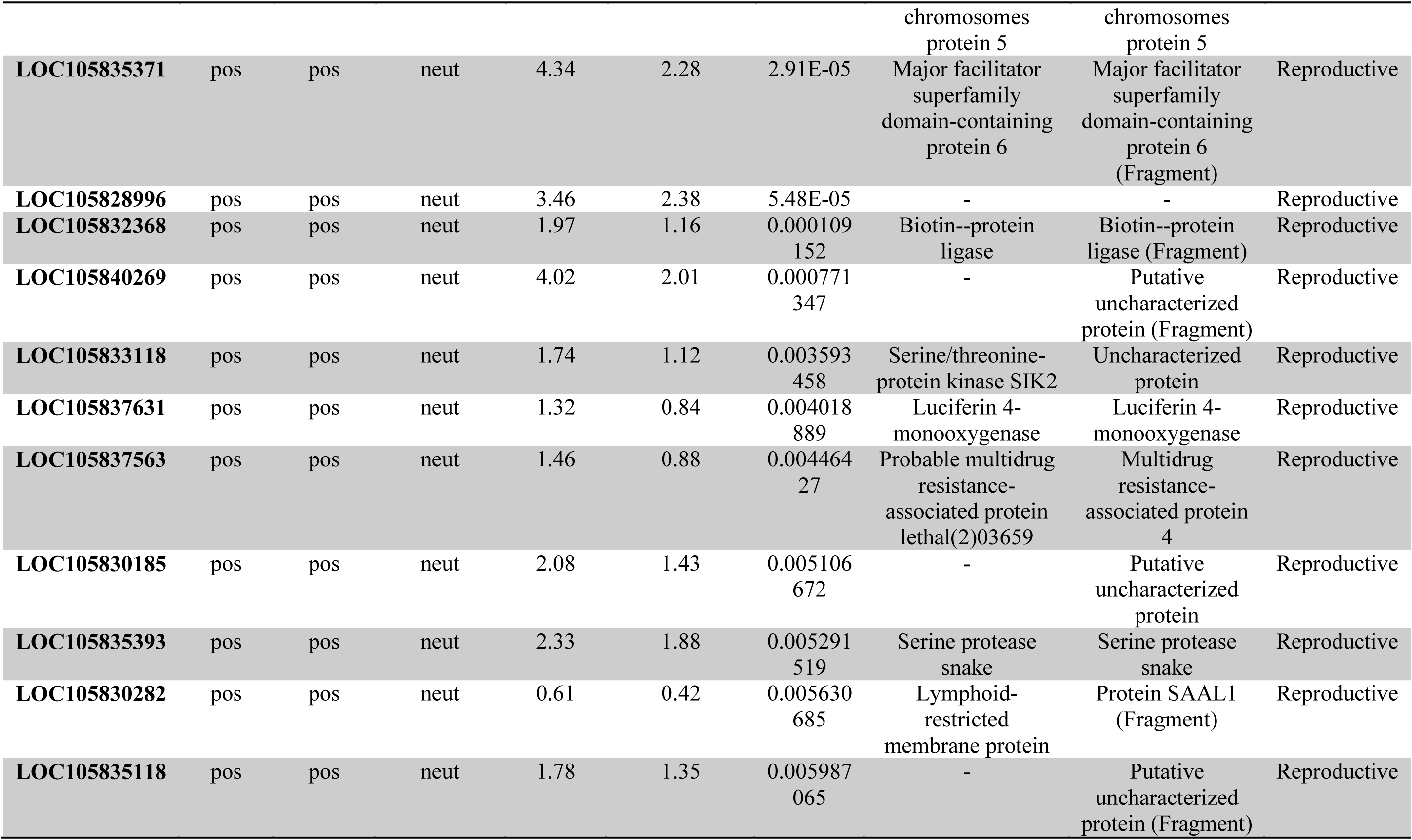
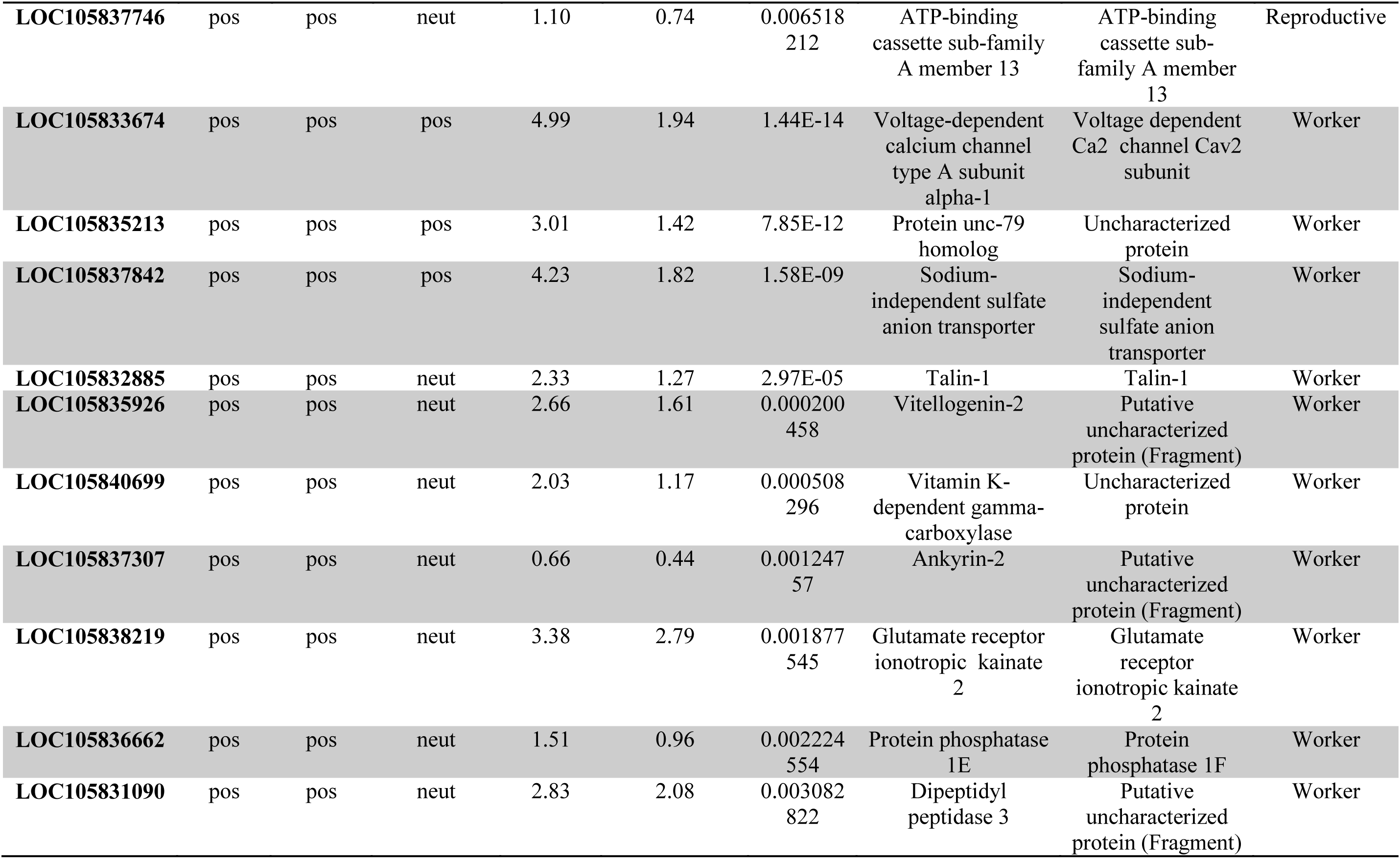
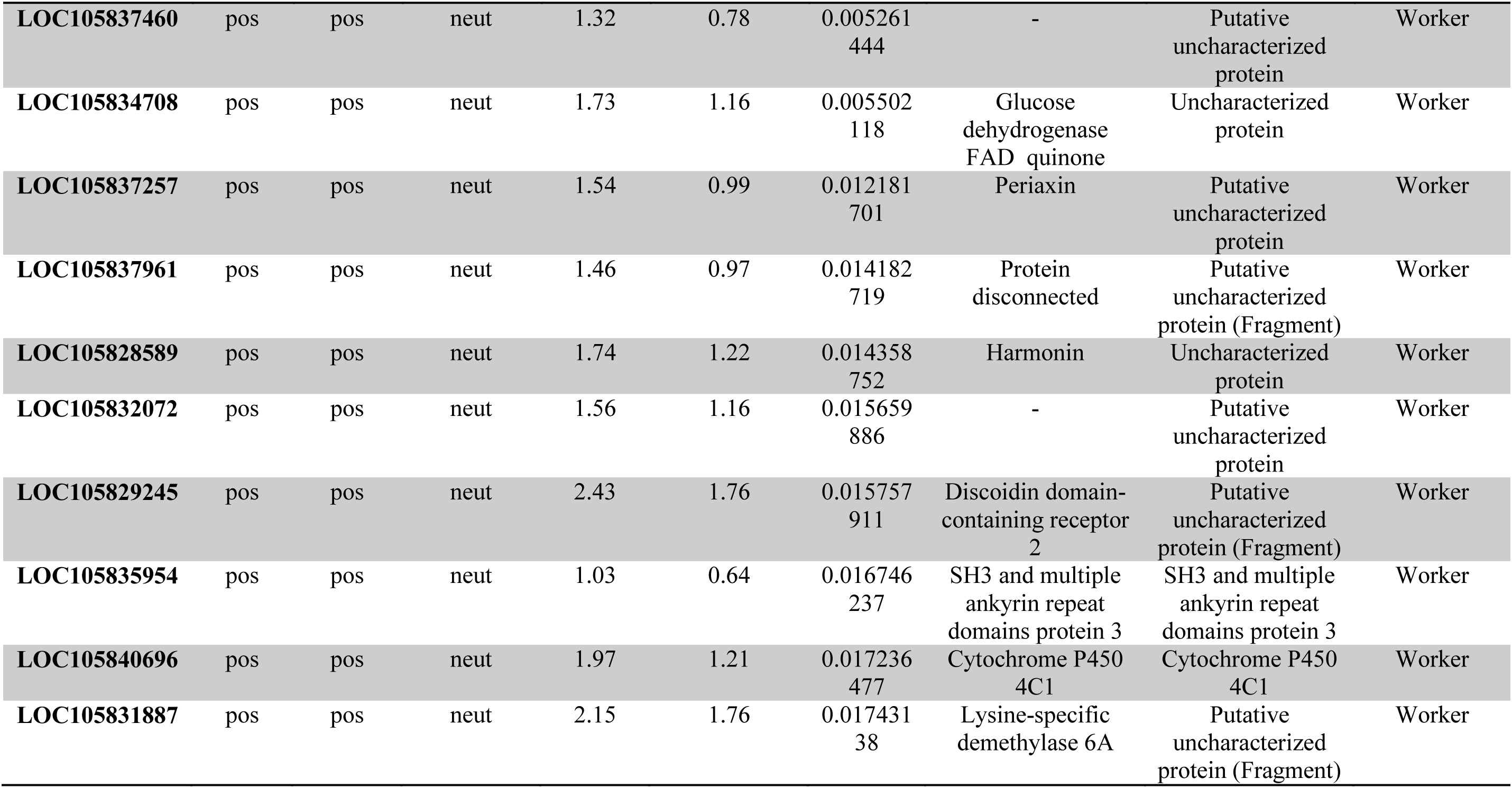
Top 20 positively selected genes (sorted by p-value of McDonald-Kreitman test) for reproductive- and worker-associated genes. SnIPRE.class is the selection categories as calculated by SnIPRE. “NI.class” refers to selection categories, as calculated using a combination of the neutrality index and the P-value from the McDonald-Kreitman test. Genes with negative values of -log10(Neutrality Index) and p-values less than 0.05 are defined as under purifying selection, while such genes with positive -log10(Neutrality Index) values are assigned to the positive selection category. “NI.class B-F correction” uses the same method but the p-value cutoff from the McDonald-Kreitman test is adjusted for multiple comparisons using the Bonferroni procedure.

### External Database S1

Complete list of genes summarizing the per-locus results of differential expression analyses, population genomic analyses, and phylostratigraphy analyses. Columns show: annotation from SwissProt and UniProt; results from differential expression analysis by larval stage (L2-L5), adult head and gaster (abdominal) tissue, across all larval samples, and across all samples, with levels NDE = non differentially expressed genes, Reproductive = reproductive-upregulated, and Worker = worker-upregulated; counts of nonsynonymous and synonymous polymorphisms within *M. pharaonis* and fixed differences between *M. pharaonis* and *M. chinense*, and total numbers of nonsynonymous and synonymous sites; results from SnIPRE analysis including BSnIPRE.class, whether genes are categorized by SnIPRE as experiencing positive selection (“pos”), negative selection (“neg”), or neither (“neut”), BSnIPRE.gamma, a population-size calibrated selection coefficient estimate, and BSnIPRE.est, normalized BSnIPRE.gamma; “NI.class” refers to selection categories, as calculated using a combination of the neutrality index and the p-value from the McDonald-Kreitman test: Genes with negative values of -log10(Neutrality Index) and p-values less than 0.05 are defined as under purifying selection, while such genes with positive -log10(Neutrality Index) values are assigned to the positive selection category. “NI.class B-F correction” uses the same method but the p-value cutoff from the McDonald-Kreitman is adjusted for multiple comparisons using the Bonferroni procedure; Finally, the assigned raw (“Raw PS”) and condensed phylostrata (“PS1” and “PS2”; Table S4) from the phylostratigraphy analyses are shown.

### External Database S2

Complete GO enrichment analysis results for workers and reproductives for each differential expression test, as calculated using the R package GOstats, sorted by p-value. L2 not included due to paucity of differentially expressed genes.

### External Database S3

Complete GO enrichment analysis results for each phylostrata category for each differential expression test, as calculated using the R package GOstats, sorted by p-value. L2 not included due to paucity of differentially expressed genes. Missing phylostrata categories returned no significant GO terms.

### External Database S4

Raw counts per locus from RNA sequencing showing level of expression across all samples included in the study (Table S1).

### External Database S5

Raw FPKM per locus from RNA sequencing showing level of expression across all samples included in the study (Table S1).

